# Genetic mapping reveals *Pou2af2*-dependent tuning of tuft cell differentiation and intestinal type 2 immunity

**DOI:** 10.1101/2022.10.19.512785

**Authors:** Marija S. Nadjsombati, Natalie Niepoth, Lily M. Webeck, Elizabeth A. Kennedy, Danielle L. Jones, Megan T. Baldridge, Andres Bendesky, Jakob von Moltke

## Abstract

Chemosensory epithelial tuft cells contribute to innate immunity at barrier surfaces, but their differentiation from epithelial progenitors is not well understood. Here we exploited differences between inbred mouse strains to identify an epithelium-intrinsic mechanism that regulates tuft cell differentiation and tunes innate type 2 immunity in the small intestine. Balb/cJ (Balb) mice had fewer intestinal tuft cells than C57BL/6J (B6) mice and failed to respond to the tuft cell ligand succinate. A majority of this differential succinate response was determined by a single genetic locus from 50-67Mb on chromosome 9 (Chr9). Congenic Balb mice carrying the B6 Chr9 locus had elevated baseline numbers of tuft cells and responded to succinate. The Chr9 locus includes *Pou2af2*, a transcriptional cofactor essential for tuft cell development. Epithelial crypts expressed a previously unannotated short isoform of *Pou2af2* that uses a novel transcriptional start site and encodes a non-functional protein. Low tuft cell numbers and the resulting lack of succinate response in Balb mice was explained by a preferential expression of the short isoform. Physiologically, differential *Pou2af2* isoform usage tuned innate type 2 immunity in the small intestine. Balb mice maintained responsiveness to helminth pathogens while ignoring commensal *Tritrichomonas* protists and reducing norovirus burdens.

**One Sentence Summary:** Genetic mapping identifies *Pou2af2* isoform usage as a novel regulator of tuft cell differentiation that tunes intestinal innate type 2 immunity.

## Introduction

Tuft cells are rare chemosensory epithelial cells that are activated by apical environmental cues and transmit signals to neighboring epithelial cells and the underlying tissue. They can be found in most mucosal barriers of mice and humans, including those of the upper airways, stomach, biliary tree, intestines, and urethra*(1)*. There are also tuft cells in the medullary thymic epithelium*(2, 3)*. Tuft cells in all tissues are defined by an apical “tuft” of long microvilli and share a transcriptional signature that includes genes also required for taste transduction (*Trpm5, Plcb2, Gnat3*) and genes associated with effector functions (*Alox5, Chat, Il25, Ptgs1*)*(4, 5)*. Although tuft cells were discovered more than 60 years ago, it wasn’t until the last ten years that studies uncovered their contributions to innate immunity at barrier tissues.

The function and ontogeny of tuft cells are perhaps best understood in the small intestine (SI), where they are one of five post-mitotic lineages of epithelial cells that comprise the SI lining. SI epithelial cells are replenished every 4-5 days from a pool of intestinal stem cells (ISCs) that reside in the crypts of the SI *(6)*. The cell intrinsic signals that direct tuft cell differentiation remain poorly understood, although the transcription factor POU2F3 is absolutely necessary for tuft cell differentiation and dispensable for all other epithelial lineages *(7)*. Progenitor cells also integrate complex environmental cues that influence their differentiation into each of the five epithelial cell types. The homeostatic cues that direct tuft cell differentiation are unknown, but in mice free of specific pathogens including *Tritrichomonas* protists, tuft cells are rare (~1% of the epithelium). On the other hand, in mice infected with helminths or colonized with *Tritrichomonas* protists, interleukin 13 (IL-13) signaling through IL-4RA and STAT6 in epithelial progenitors is necessary and sufficient to induce a 5-10 fold increase in tuft cell frequency (hyperplasia) *(7–10)*.

SI tuft cells detect the presence of helminths and *Tritrichomonas* protists using a signaling pathway related to taste transduction in the tongue *(9)*. The ligand and receptor required for helminth sensing remain unknown, but Tritrichomonads are sensed via secretion of the metabolite succinate, which binds to its receptor (SUCNR1) on tuft cells*(5, 11, 12)*. Once activated, tuft cells produce the cytokine interleukin 25 (IL-25) and, in some contexts, lipid-derived cysteinyl leukotrienes (cysLTs) to directly activate group 2 innate lymphoid cells (ILC2s) in the SI lamina propria (SILP)*(7–9, 13)*. ILC2s in turn produce the canonical type 2 cytokines IL-5, −9, and −13, which collectively regulate hallmarks of type 2 immunity, including eosinophilia, hyperresponsivity of smooth muscle, and mucus overproduction. Meanwhile, the IL-13-induced tuft cell hyperplasia establishes a feed-forward tuft-ILC2 circuit. Tuft-ILC2 circuit activation can promote helminth clearance, but the function of tuft cell hyperplasia during *Tritrichomonas* colonization remains unclear as these protists are acquired from parents post-partum and persist for the life of the mouse*(7–9)*.

The tuft-ILC2 circuit is regulated differently in the proximal (i.e. duodenum + proximal jejunum) and distal (i.e. distal jejunum + ileum) ends of the SI. Helminths that primarily reside in the proximal SI, such as *Nippostrongylus brasiliensis (Nb)* and *Heligmosomoides polygyrus* (*Hp*), induce both IL-25 and cysLTs to elicit stronger tuft-ILC2 activation in the proximal SI compared to the distal SI*(13)*. Conversely, succinate activation of the tuft-ILC2 circuit is cysLT-independent and occurs most strongly in the distal SI, where *Sucnr1* expression is highest and protists are more abundant*(12, 13)*. Even when succinate is provided in the drinking water, and therefore is at its highest concentration in the proximal SI, tuft cell hyperplasia predominates in the distal SI*(5)*.

Studying different inbred strains of mice has provided insight into mechanisms of type 2 immunity. In particular, seminal studies using *Leishmania major* infection established the notion that CD4^+^ T helper cell responses in Balb/cJ (Balb) mice are type 2 biased, while C57BL/6J (B6) mice are type 1 biased*(14, 15)*. Such biases have similarly been noted in the SI, where Balb mice clear some helminth infections, such as *Hp* and *Strongyloides ratti*, more efficiently than B6 mice*(16, 17)*. There are many proposed mechanisms for these differences, including intestinal microbiota composition, regulatory T cell function, and strength of T helper 2 cell (Th2) activation*(16, 18, 19)*. Some studies have used genetic mapping to identify loci associated with strain-specific immune responses, but no single gene is responsible for the observed phenotypes*(20, 21)*. Likely these differences arise from a complex network of genetic and environmental differences all contributing to the phenotypic outcomes. Whether tuft cells and the SI tuft-ILC2 circuit are differentially regulated across mouse strains has not been examined.

Here we find that Balb mice have fewer tuft cells than B6 mice in many tissues and fail to activate the tuft-ILC2 circuit following succinate treatment in the SI. These differences are determined by a single genetic locus that regulates tuft cell differentiation and tunes the sensitivity and kinetics of innate type 2 immunity in the SI.

## Results

### Balb mice have fewer intestinal tuft cells and do not respond to succinate

Given that Balb mice have been described as “type 2 skewed”, but nearly all tuft cell studies have been performed in B6 mice, we set out to compare tuft cell frequency and function between B6 and Balb mice. Balb mice were previously reported to have fewer tuft cells than B6 in the distal SI*(22)*, and we found this discrepancy extended throughout the intestinal tract (Fig. 1A–B, fig. S1A). Balb mice also had a trend towards fewer tuft cells in the trachea, but equivalent frequencies of tuft cells in the thymus by flow cytometry (Fig. 1A–C).

**Figure 1.**
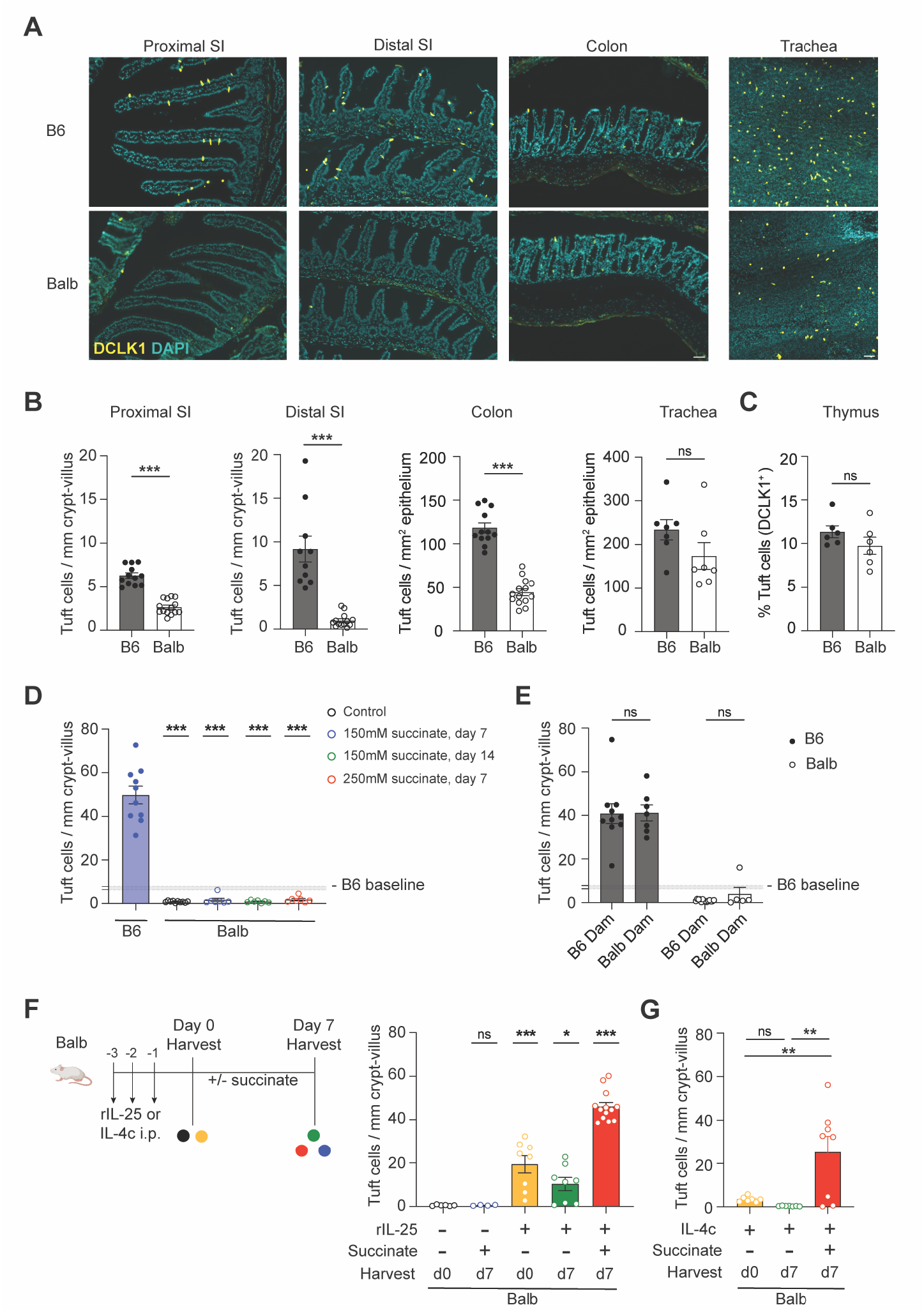
Balb mice have fewer tuft cells at baseline and do not develop succinate induced hyperplasia unless primed. (**A** and **B**) (A) Representative images and (B) tuft cell (DCLK1+) quantification by immunofluorescence from indicated tissues and indicated mice. (**C**) Thymic tuft cell quantification by flow cytometry. (**D**) Tuft cell quantification in the distal SI of Balb mice at indicated succinate concentrations and time points. (**E**) Tuft cell quantification in the distal SI of adult B6 and Balb mice raised by dams of indicated genotype and given 150mM succinate for 7 days. (**F-G**) Experimental schematic and tuft cell quantification in the distal SI of Balb mice treated with either (F) rIL-25 or (G) IL-4c as indicated. In the graphs, each symbol represents an individual mouse from three or more pooled experiments. In (D and E), shaded area indicates the 95% confidence interval of the mean for distal SI tuft cell quantification calculated from a large cohort of control B6 mice. *p < 0.05, **p < 0.01, ***p < 0.001 by Mann-Whitney (B and C), by one way ANOVA with comparison to B6 (D) or Balb untreated (G) or multiple comparisons (H), and by multiple t tests (E). n.s., not significant. Graphs depict mean +/− SEM. Also see Figure S1.

Next, we tested the tuft-ILC2 circuit in Balb mice. As previously described*(5, 11, 12)*, B6 mice given 150mM sodium succinate in the drinking water for 7 days developed robust tuft cell hyperplasia in the distal SI (Fig. 1D). Balb mice given succinate drinking water failed to induce tuft cell hyperplasia, even if succinate was administered for longer (14 days) or at a higher dose (250mM for 7 days). We tested whether the defect in succinate-induced hyperplasia in Balb mice was microbiome dependent by cross-fostering Balb and B6 litters. At adulthood, Balb mice raised by a B6 dam still failed to develop tuft cell hyperplasia following succinate administration (Fig. 1E). In fact, even after succinate administration, the tuft cell frequency in Balb mice raised by B6 dams remained below the baseline B6 level. Conversely, B6 mice developed hyperplasia regardless of dam. We therefore conclude that the microbiome is not responsible for the homeostatic and induced frequency of tuft cells in B6 and Balb mice.

To test whether responses to succinate are restored when the starting tuft cell number in Balb mice is elevated, we ‘primed’ the tuft-ILC2 circuit of Balb mice by giving recombinant IL-25 (rIL-25) to directly induce IL-13 release from ILC2s and increase tuft cell frequency (Fig. 1F). When rIL-25 treatment was followed by a week of regular drinking water, tuft cell frequency returned towards baseline. However, mice pre-treated with rIL-25 and then given succinate for 7 days readily developed tuft cell hyperplasia. Similar results were achieved if IL-4 complex (IL-4c), which recapitulates the signaling effects of IL-13 on stem cells, was administered to increase tuft cell frequency prior to succinate administration (Fig. 1G).

Although we have never noted sex-dependent differences in tuft cell frequency or succinate responsiveness in B6 mice (data not shown), some male Balb mice given IL-4c still failed to respond to succinate (fig. S1B). The effect of sex on tuft cells themselves is unknown, but studies of airway ILC2s have demonstrated androgen-dependent reductions in ILC2 activation*(23–27)*. We hypothesize that SILP ILC2s are similarly impacted, but that this is only revealed in “sensitized” contexts where ILC2s are weakly activated. Nonetheless, these priming experiments demonstrate that the tuft-ILC2 circuit is intact but hyporesponsive in Balb mice.

### ILC2s are abundant and functional in Balb mice

The tuft-ILC2 circuit contains three cellular components: mature tuft cells, ILC2s, and epithelial stem cells. To determine which component accounts for the Balb defect, we began by assessing the number, phenotype and cytokine production of ILC2s. Compared to B6 mice, unmanipulated Balb mice actually had more ILC2s (CD45^+^, Lin^−^, GATA3^+^) in the distal SILP (Fig. 2A–B, fig. S2A). The expression of the IL-25 receptor subunit IL-17RB was equivalent between the two strains, while CD44 and KLRG1, markers of lymphocyte activation, were both reduced on Balb ILC2s (Fig. 2C–E). We previously noted similarly-reduced KLRG1 expression on SILP ILC2s from unmanipulated *Tritrichomonas-*free B6.*Il25*^-/-^ and B6.*Trpm5*^-/-^ mice, suggesting tonic signaling from tuft cells to ILC2s in the absence of known tuft cell ligands *(13)*. Therefore, by analogy to these mice, the low frequency of tuft cells in Balb mice likely leads to loss of tonic signaling and accounts for the lower KLRG1 expression. As before, and consistent with studies in the lung*(27)*, we noted sex-dependent differences in KLRG1, but in all cases the Balb ILC2s had lower KLRG1 expression than sex-matched B6 ILC2s (Fig. 2E). Male ILC2s in the SILP have higher expression of KLRG1, yet develop less robust tuft cell hyperplasia in some assays (Fig. 1G, fig. S1B). While counterintuitive, both findings are consistent with studies in the lung, where male ILC2s have higher KLRG1 on a population level, yet are less activated following stimulation*(23, 27)*. Finally, there was no difference in the number of GATA3^+^ Th2 cells or eosinophils in the SILP of Balb and B6 mice (Fig. S2A–C).

**Figure 2.**
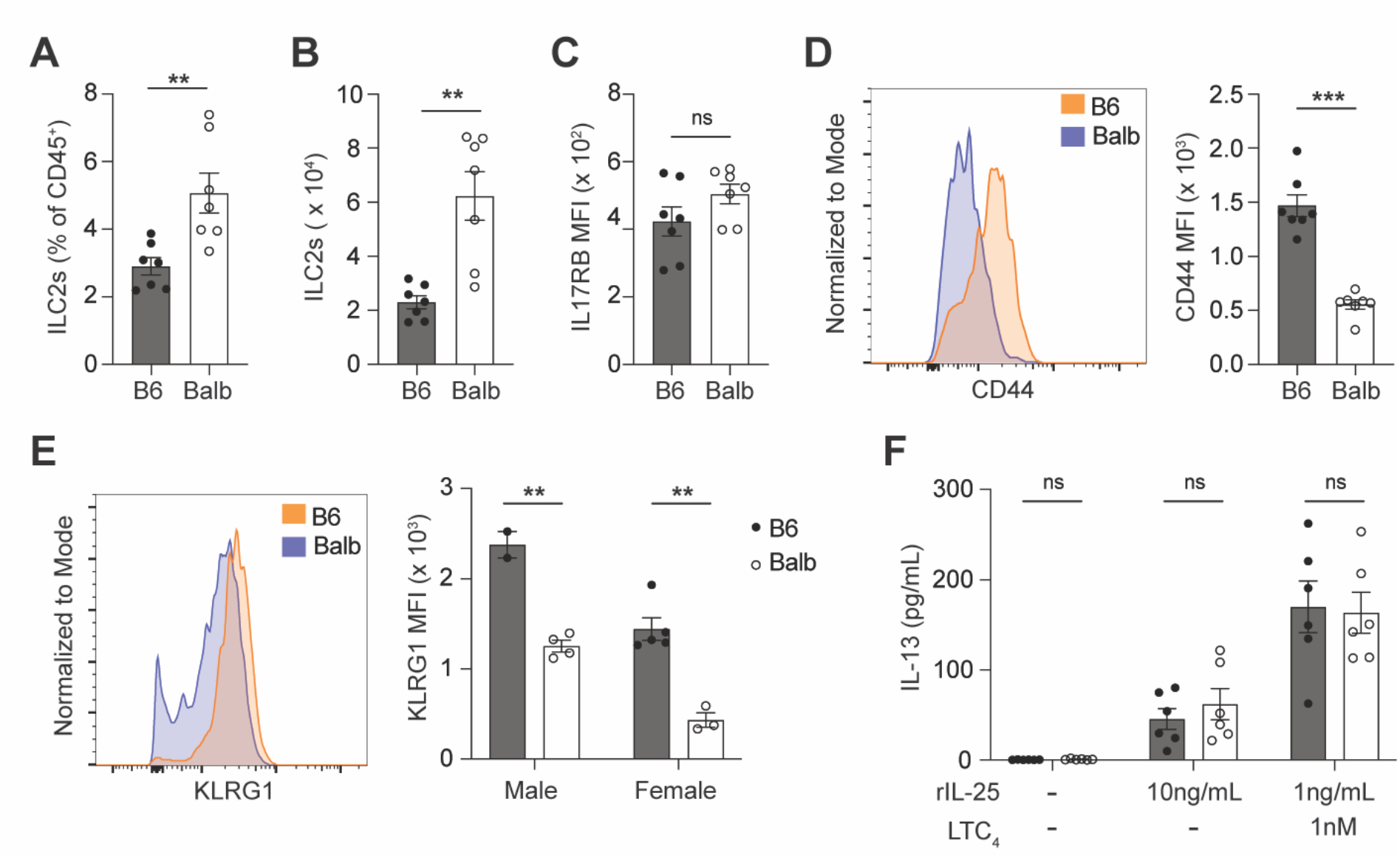
Balb ILC2s are equally responsive to IL-25 but less activated at baseline compared to B6 ILC2s. (**A** and **B**) Quantification of ILC2s (CD45^+^ Lin^−^ GATA3^+^) by (A) percentage and (B) absolute number in the SILP. (**C**, **D** and **E**) Quantification of (C) IL17RB MFI (D) CD44 MFI and (E) KLRG1 MFI on ILC2s. (**F**) IL-13 concentration in the supernatant following 6-h *in vitro* culture of SI ILC2s with the indicated concentrations of rIL-25 and LTC_4_. In the graphs, each symbol represents an individual mouse from two pooled experiments. *p < 0.05, **p < 0.01, ***p < 0.001 by Mann-Whitney (A –D) or by multiple t tests (E and F). n.s., not significant. Graphs depict mean +/− SEM. Also see Figure S2.

To test the functional capacity of Balb ILC2s, we sorted ILC2s from the SILP of unmanipulated mice and cultured them with rIL-25 with or without leukotriene C_4_ (LTC_4_). B6 and Balb ILC2s made moderate but equivalent amounts of IL-5 and IL-13 following 6 hours of rIL-25 treatment (Fig. 2F, fig. S2D). Cytokine production was greatly enhanced by the addition of LTC_4_, but IL-13 and IL-5 secretion was still equivalent between the two strains. Additionally, B6 and Balb ILC2s had equivalent expression of the proliferation marker Ki67 two days post stimulation (fig. S2E). No sex differences were observed in this assay. Overall, compared to B6, Balb ILC2s are more abundant and equally capable of responding to tuft cell signals in the SILP. The failure of Balb mice to activate the tuft-ILC2 circuit in response to succinate is therefore likely ILC2-independent.

### The Balb tuft cell defect is epithelium intrinsic

Next, we used organoids to determine whether the Balb tuft cell defect was epithelium intrinsic or required signals from surrounding stromal or immune cells. Organoid cultures contain only epithelial cells and recapitulate epithelial differentiation, including IL-13-induced tuft cell hyperplasia*(7–9)*. Tuft cell frequency was significantly lower in untreated Balb organoids compared to B6 organoids after both 1 and 4 weeks in culture, demonstrating that the tuft cell defect is epithelium intrinsic and stably maintained *ex vivo* (Fig. 3A–B, Fig. S3A). Recombinant IL-13 (rIL-13) induced tuft cell hyperplasia in organoids from both strains; however, Balb organoids had a lower frequency of tuft cells when compared to B6 organoids, particularly when cultured 4 weeks before rIL-13 treatment (Fig. 3C). Given that Balb organoids started from a lower baseline, the fold increase in tuft cells induced by rIL-13 was greater in Balb organoids, suggesting their defect predominantly impacts IL-13-independent tuft cell differentiation (Fig. S3B).

**Figure 3.**
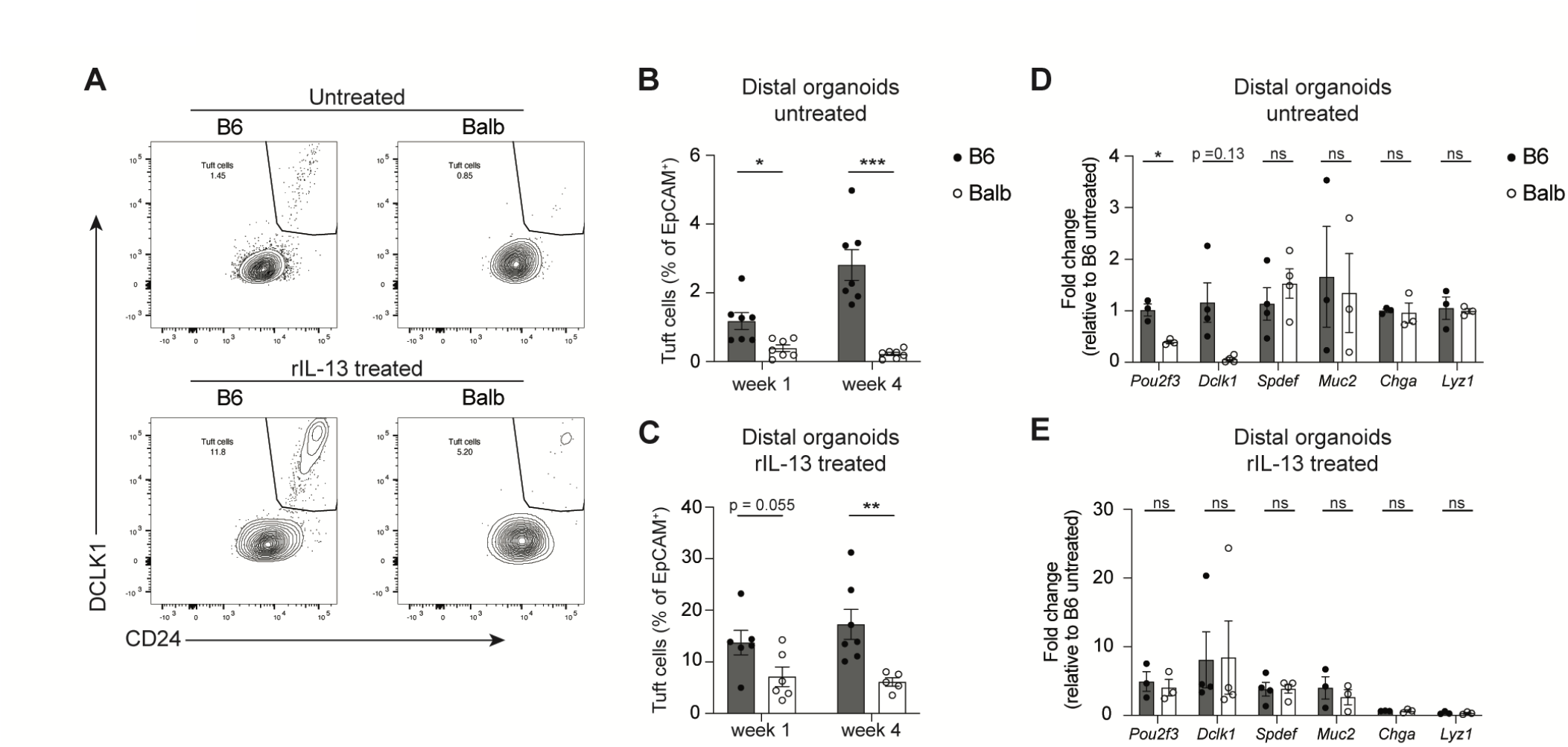
Balb tuft cell defect is epithelium intrinsic and tuft cell specific. (**A**) Representative flow cytometry plots of tuft cell quantification from B6 or Balb distal SI organoids cultured *in vitro* for one week, either untreated or rIL-13 treated (2.5 ng/ml). (**B** and **C**) Quantification of tuft cells from (A) (**D** and **E**) Real-time PCR quantification of indicated genes normalized to B6 untreated condition, all relative to *Rps17* expression from (D) control or (E) rIL-13 treated distal SI organoids cultured for 2 weeks *in vitro*. In the graphs, each symbol represents a biological replicate based on the average of 2 to 3 technical replicates, from three to six pooled experiments. *p < 0.05, **p < 0.01, ***p < 0.001 by multiple t tests (B - E). n.s., not significant. Graphs depict mean +/− SEM. Also see Figure S3.

To assess if the Balb defect is specific to tuft cells, we used qPCR to quantify all secretory cell lineages in organoids after 2 weeks in culture. Untreated Balb organoids had significantly lower expression of the tuft cell gene *Pou2f3*, and a strong trend toward less *Dclk1* (Fig. 3D). Expression of the goblet cell genes *Spdef* and *Muc2*, the Paneth cell marker *Lyz1* and the enteroendocrine cell marker *Chga* was equivalent between Balb and B6 organoids (Fig. 3D). Using this transcriptional measure, there was no difference between rIL-13 treated Balb and B6 organoids for any gene, including tuft cell markers (Fig. 3E). The discrepancies with Fig. 3C suggest post-transcriptional regulation of tuft cell differentiation, but also further support the conclusion that the Balb defect predominantly impacts homeostatic tuft cell differentiation. Together, these data demonstrate a tuft cell-specific and epithelium-intrinsic reduction in Balb mice.

### Differential tuft cell frequency between B6 and Balb mice is genetically regulated

We next hypothesized that the Balb and B6 differences in tuft cell frequency and succinate responsiveness are determined by genetic differences between the two strains. To test heritability of succinate responsiveness, we generated Balb x B6 F1 and F2 mice and measured tuft cell frequency after succinate treatment. In this context we again noted reduced succinate responses in male mice (fig. S4A–B), so we assessed succinate response rates using female F1 and F2 mice. F1 mice all developed tuft cell hyperplasia when treated with succinate, but there were both responsive and nonresponsive mice in the F2 generation (Fig. 4A). Using a cutoff of 13 tuft cells/mm crypt villus, which is just above the B6 baseline, 80% of F2 female mice responded to succinate. The frequency of succinate responsive F1 and F2 mice is therefore consistent with a single recessive locus determining tuft cell differences between Balb and B6 mice.

**Figure 4.**
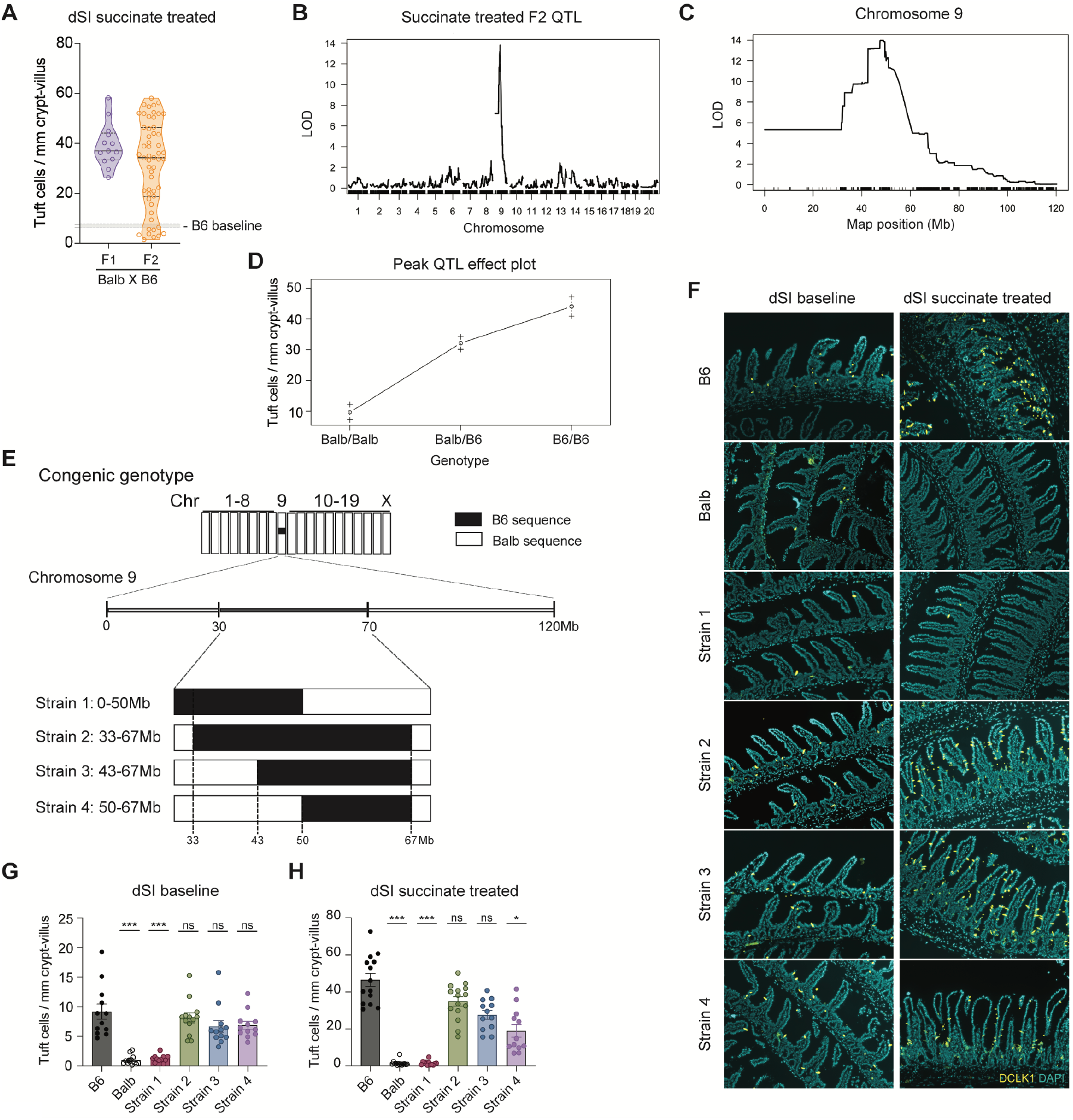
A single locus on chromosome 9 regulates baseline tuft cell frequency and succinate responsiveness. (**A**) Quantification of tuft cells from distal SI of succinate treated female mice. (**B** and **C**) QTL mapping of succinate induced tuft cell hyperplasia in Balb X B6 F2 cross (B) whole genome and (C) zoomed in on Chr9. (**D**) Effect plot of tuft cell phenotype based on genotype at the peak QTL (Chr9:50857809) (**E**) Schematic of genotype for congenic Strain 1-4 mice. (**F - H**) (F) Representative images and quantification of tuft cells from distal SI at (G) baseline or (H) after 150mM succinate treatment. Some B6 and Balb data points shown in (G) and (H) are also included as controls in Figure 1B and 1D. In (A), shaded area indicates the 95% confidence interval of the mean for distal SI tuft cell quantification calculated from a large cohort of control B6 mice. In the graphs, each symbol represents an individual mouse from three or more pooled experiments. *p < 0.05, **p < 0.01, ***p < 0.001 by one-way ANOVA (G and H) with comparison to B6. n.s., not significant. Graphs depict mean +/− SEM. Also see Figure S4.

Combining high-density single nucleotide polymorphism (SNP) genotyping with phenotypes of F2 mice is a powerful technique for identification of quantitative trait loci (QTL) that explain phenotypic variation. We therefore employed low-coverage whole genome sequencing coupled to imputation to genotype B6 x Balb F2 mice *(28)*. In brief, Tn5 transposase was used to randomly insert DNA tags into genomic DNA from 84 B6 x Balb F2 mice, here analyzing both male and female mice. Tag-adjacent genomic sequences were obtained by next-generation sequencing and assigned to B6 and Balb genomes to provide whole-genome genotyping at higher resolution than traditional SNP-based arrays. We then combined these genotypes with the succinate-induced tuft cell frequency of each F2 mouse and performed QTL mapping. We detected a dominant QTL on chromosome 9 (Chr9), with a peak at 50,857,809 bp and 1.5 LOD support interval from 45.49 Mb to 53.03 Mb (Fig. 4B–C). Sex was a significant additive covariate, but the Chr9 locus was dominant in both sexes (fig. S4C). An effect plot at the peak QTL location revealed a clear gene dosage dependent response to succinate (Fig. 4D), and the Chr9 locus explained 53% of the variance in succinate response in this 84-mouse F2 cohort. Thus, a single locus accounts for a majority of the difference in succinate-induced tuft cells between B6 and Balb mice.

To begin fine mapping and to generate congenic mice in which only the Chr9 locus is B6-derived, we initiated a series of backcrosses. B6 x Balb F1 mice were crossed to wild-type Balb mice and resulting offspring were again crossed to wild-type Balb mice for 6 to 8 generations. In each generation, we used low-coverage whole genome sequencing coupled to imputation to look for crossover events that reduced the size of the Chr9 locus and to identify mice that had lost B6 DNA in other regions (a process sometimes called speed congenics). This process generated 4 strains of congenic mice carrying distinct B6-derived portions of the Chr9 QTL and homozygous for Balb DNA at all other locations (Fig. 4E).

Congenic strains 2-4 had B6-equivalent levels of tuft cells in the distal SI at baseline and developed tuft cell hyperplasia when succinate treated (Fig. 4F–H). Tuft cell frequency was also increased in proximal SI, colon, and trachea of Strain 3 mice compared to Balb (Fig. S4D–F). Strain 1 had Balb-equivalent levels of tuft cells at baseline and did not develop tuft cell hyperplasia after succinate treatment (Fig. 4F–H). Strain 1 contains B6 sequence from 33-50Mb, indicating the relevant locus in not within this region. Instead, the locus controlling baseline tuft cell number and succinate responsiveness is in the 50-67Mb region shared by Strains 2, 3, and 4. In sum, the 50-67 Mb region of Chr9 explains most of the differential succinate sensing in B6 and Balb mice, and when placed in a Balb genome restores B6 levels of homeostatic tuft cells and succinate responsiveness in the distal SI.

### RNA sequencing identifies *1810046K07Rik / Pou2af2* as a gene of interest

Genetic variation within species often shapes traits by either changing the expression of genes or the amino acid sequence of the proteins they encode. To discover the gene(s) mediating the difference, we systematically compared the congenic interval (Chr9:50-67 Mb) in B6 and Balb mice. We found that the interval contains ~3170 genetic variants that distinguish B6 and Balb genomes. Many of these variants are in intergenic regions or associated with genes that are not expressed in the SI epithelium, but even focusing on the 1.5 LOD confidence interval and on variants predicted to alter protein sequence, it was difficult to identify candidates for further investigation.

To assess gene expression and leverage the congenic strains, we sequenced the mRNA of tuft cells (CD45^−^ EPCAM^+^ SigF^+^ CD24^+^) sorted from the distal SI of B6, Balb, and congenic Strain 3 mice and identified differentially expressed genes (DEGs; log2FC > 1, FDR < .05) (Fig. S5A–C). Hierarchical clustering of all DEGs revealed 3 expression modules (Fig. 5A). Within each module, we looked for genes that were part of the SI tuft cell signature and/or located in the Chr9 locus *(5)* (Table S1). Module 1 was comprised of genes more highly expressed in Balb and congenic than B6. One gene, *Hebp1*, was a tuft cell signature gene, but is not located on Chr9. *Gm7293*, encoded at 51.5 Mb on Chr9 was also in this module. Module 2 contained a subset of 31 genes enriched selectively in Balb samples. Many of these genes were Paneth cell related, such as lysozyme and defensins, and likely represented low-level contamination by CD24^+^ Paneth cells, which is amplified in Balb samples due to the rarity of tuft cells in these mice. None of these genes are located on Chr9. Module 3 contained genes more highly expressed by B6 tuft cells compared to congenic or Balb. There were several tuft cell signature genes within this module, including *Sucnr1* (Fig. 5B). *Sucnr1* is encoded on Chr3, is unlikely to directly impact tuft cell differentiation, and was not upregulated in congenic Strain 3 tuft cells, so it does not explain baseline differences in tuft cell frequency. Reduced *Sucnr1* expression could, however, contribute to the failure of Balb mice to sense succinate, yet the effect of the Chr9 locus in Strain 3 mice was enough to restore succinate sensing despite Balb-equivalent *Sucnr1* expression (Fig. 4H).

**Figure 5.**
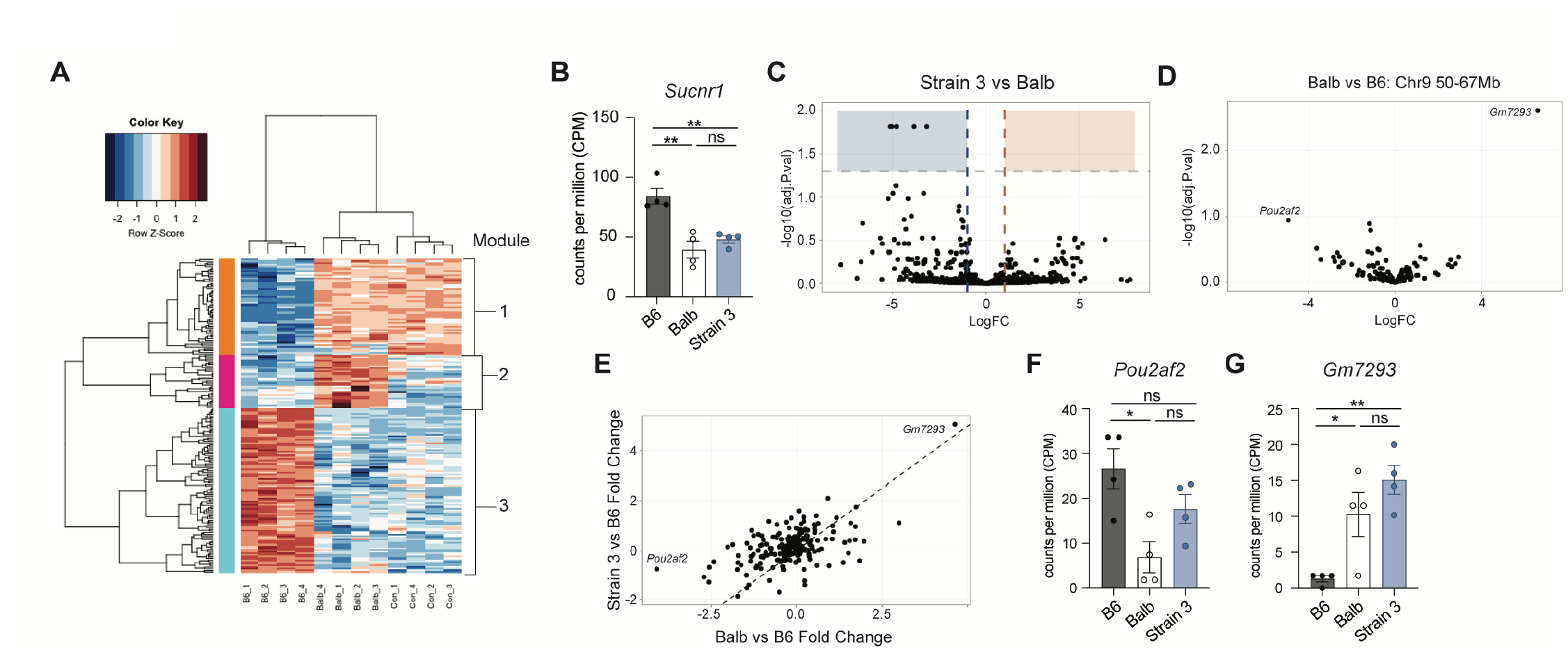
mRNA sequencing of mature tuft cells from B6, Balb and Strain 3 mice. (**A**) Hierarchical clustering of differentially expressed genes. (**B**) Normalized read count of *Sucnr1*. (**C**) Volcano plots depicting DEGs from Strain 3 vs Balb. (**D**) Volcano plots depicting DEGs for genes found in the Chr9 50-67Mb region, from Balb vs B6. (**E**) Plot of fold change of DEGs from Strain 3 vs B6 compared to fold change from Balb vs B6. (**F** and **G**) Normalized read count of (F) *Pou2af2* and (G) *Gm7293*. *p < 0.05, **p < 0.01, ***p < 0.001 by one-way ANOVA (B, F and G). n.s., not significant. Graphs depict mean +/− SEM. Also see Figure S5.

To focus on transcriptional regulation revealed by the congenic mice, we performed a pair-wise comparison between Balb and Strain 3, but identified only 5 DEGs, and none are encoded on Chr9 (Fig. 5C). Since genome-wide DEG analysis did not identify any candidate genes, we specifically compared genes from the Chr9 locus (50-67Mb) in B6 and Balb mice (Fig. 5D). *Gm7293* again appeared as highly upregulated in Balb mice, while *1810046K07Rik* was the top downregulated gene. *1810046K07Rik* also stood out as the only tuft cell signature gene that was downregulated in Balb mice but rescued in Strain 3 congenic mice, while *Gm7293* expression was unchanged between Balb and Strain 3 (Fig. 5E–G).

*1810046K07Rik* and another gene, *Colca2*, were recently found to encode co-factors required for the function of POU2F3*(29, 30)*. These genes, and the proteins they encode, were respectively renamed *Pou2af2/*OCA-T1 and *Pou2af3/*OCA-T2 and are located in a gene cluster together with *Pou2af1/*OCA-B. OCA-B is a co-factor for OCT-1 and OCT-2, transcription factors closely related to POU2F3*(31)*. This gene cluster is located at 51.2 Mb on Chr9, very close to the QTL peak (50.8 Mb).

### Differential *Pou2af2* isoform expression in intestinal crypts

*Pou2af2^-/-^* mice were reported to lack tuft cells in the SI and trachea, but have normal tuft cell numbers in the thymus, a distribution similar to our findings in Balb mice*(29)*. *Pou2af3^-/-^* mice have not yet been generated, but *Pou2af3* expression is low or undetectable in RNA sequencing of SI tuft cells, and it is not included in the SI tuft cell signature*(5)*. We therefore focused our attention on identifying a mechanism by which *Pou2af2* might regulate differential tuft cell phenotypes in B6 and Balb mice.

Because *Pou2af2* is currently annotated with two transcriptional start sites, we used 5’ Rapid Amplification of cDNA Ends (RACE) for unbiased amplification of *Pou2af2* transcripts. Since the few mature tuft cells that emerge in Balb mice may not represent events that occur during differentiation, we used RNA from distal SI crypts to capture tuft cell progenitor cells. A primer designed to capture all annotated isoforms produced ~550bp and ~450bp bands in B6 samples. Balb samples lacked the 550bp band but contained the 450bp band and a faint ~100bp band (Fig. 6A). Cloning and sequencing of these bands revealed that the 100bp band resulted from non-specific amplification of 18S RNA and the 550bp band corresponded to the full-length *Pou2af2* isoform (Fig. 6B). The 450bp band present in both Balb and B6 samples, however, corresponded to an isoform not listed in the current *Mus musculus* genome release (GRCm39) and did not appear to use either of the annotated transcriptional start sites. This isoform begins 26bp downstream of the annotated transcription start site in exon 2 of the full-length isoform. The first available translation start site in this isoform gives rise to a truncated protein that lacks a 20 amino acid N-terminal motif shared by OCA-T1, OCA-T2, and OCA-B, and required for binding to their target transcription factors (i.e. POU2F3 or OCT-1/2)*(29, 32)*. We did not find evidence that any of the other annotated isoforms were expressed in SI crypts.

**Figure 6.**
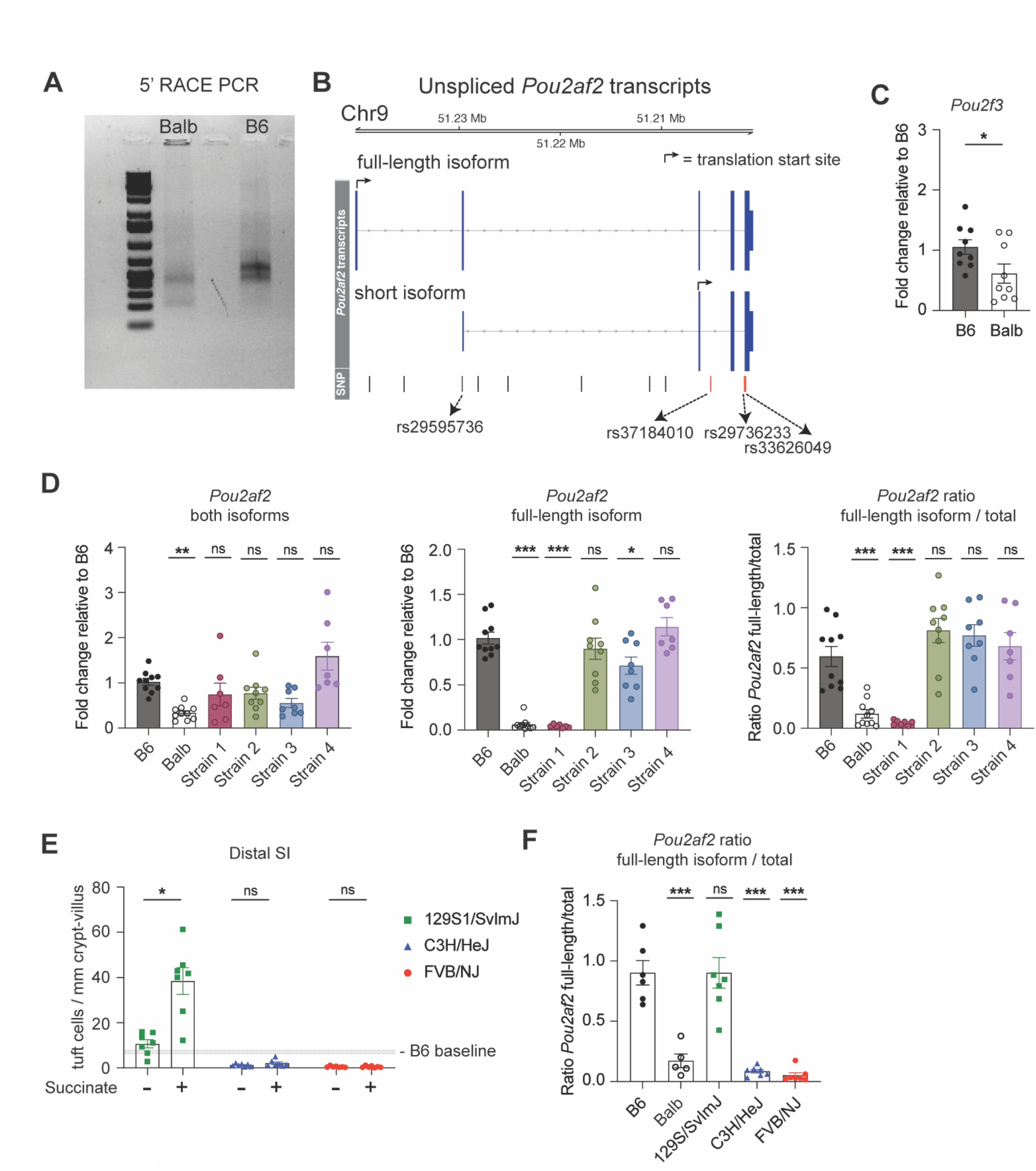
*Pou2af2* isoform expression is modulated by genotype. (**A**) Agarose gel of 5’ Rapid amplification of cDNA ends products from distal SI crypts. (**B**) Schematic of *Pou2af2* isoforms expressed in distal SI crypts with annotated SNPs (vertical bars) that differ between B6 and Balb. SNPs that also match phenotypes of other inbred strains are highlighted in red. (**C)**Real-time PCR quantification of *Pou2f3*. (**D**) Real-time PCR quantification of indicated *Pou2af2* isoform and *Pou2af2* isoform ratio. (**E**) Tuft cell quantification in the distal SI and (**F**) *Pou2af2* isoform ratio calculated from real-time PCR quantification from distal SI crypts of indicated strains. In the graphs, each symbol represents an individual mouse three or more pooled experiments. *p < 0.05, **p < 0.01, ***p < 0.001 by Mann-Whitney (C), by one-way ANOVA (D and F) with comparison to B6 and by multiple t-tests (E). n.s., not significant. Graphs depict mean +/− SEM. Also see Figure S6.

The results of the 5’ RACE allowed us to design qPCR primers to quantify the abundance of the full-length transcript only and of all transcripts combined. No primers could be designed for just the short isoform, as it shares 100% homology with the full-length isoform. Due to the lack of tuft cells in Balb mice, Balb SI crypts had significantly lower expression of *Pou2f3* and both isoforms of *Pou2af2* than B6 crypts (Fig. 6C–D). Within each sample, however, the portion of total *Pou2af2* transcript accounted for by full-length transcript is also significantly lower in Balb crypts compared to B6 (Fig. 6D). Crypts from congenic strain 1 phenocopy Balb crypts with about 10% of total *Pou2af2* transcript being the full-length isoform. Congenic strains 2,3 and 4 express 70-80% full-length isoform, similar to B6 (Fig. 6D).

We also used qPCR to analyze *Pou2af3* expression and isoform usage. *Pou2af3* has a full-length and a short isoform that each contain unique portions, allowing us to design isoform-specific primers. As with *Pou2af2*, the short isoform of *Pou2af3* lacks the POU2F3 binding domain. As expected, SI crypt expression of *Pou2af3* is lower than *Pou2af2* regardless of strain and lower in Balb and Strain 1 than B6 and Strain 2-4 (fig. S6A). The full-length isoform in particular was nearly undetectable in all mice. Nonetheless, Balb and Strain 1 had a decreased ratio of full-length to total *Pou2af3* and increased ratio of short isoform to total *Pou2af3* compared to B6, Strain 2, 3 and 4 (fig. S6B–C). Finally, as in the organoids, there was no difference in expression of genes for other lineages of epithelial cells between B6 and Balb crypts, confirming that the Balb defect is tuft cell specific (fig. S6D).

To understand if the differences between B6 and Balb mice were generalizable, we examined baseline tuft cells and succinate sensing in additional strains of mice. Succinate responsiveness was highly variable in Swiss Webster mice, an outbred strain, suggesting genetic diversity can lead to diverse succinate responses (Fig. S6E). Testing inbred strains, we found FVB/NJ and C3H/HeJ strains had very low numbers of tuft cells at baseline and did not develop tuft cell hyperplasia following succinate treatment, phenocopying Balb (Fig. 6E). On the other hand, 129S1/SvImJ mice had close to B6 levels of tuft cells at baseline and upon succinate treatment developed tuft cell hyperplasia in the distal SI. We measured *Pou2af2* isoform expression in distal SI crypts from these strains. The ratio of full-length isoform to total *Pou2af2* expression corresponded with baseline tuft cell number and succinate phenotype, with 129S1/SvImJ mice having a high ratio, similar to B6, and FVB/NJ and C3H/HeJ mice having a low ratio, similar to Balb (Fig. 6F). Although total *Pou2af3* expression was again lower than *Pou2af2*, the ratio of short and long isoforms followed the same trend as *Pou2af2*. (fig. S6F). It appears, therefore, that *Pou2af2* and *Pou2af3* are somehow coregulated, but given the higher expression of *Pou2af2* and the similarities between *Pou2af2^-/-^* and Balb mice, we propose that the production of fewer mature tuft cells in Balb and Strain 1 mice results from a lack of functional OCA-T1 expression and therefore a failure to induce POU2F3-dependent gene transcription.

### Analysis of genetic variants in *Pou2af2* locus

Analysis of genetic variants in the *Pou2af2* locus revealed 11 single nucleotide polymorphisms (SNPs) that distinguish B6 and Balb mice, several of which may be of interest (Fig. 6B). First is rs29595736, located in exon 2 of the full-length isoform and just upstream of the transcriptional start site of the short isoform (fig. S6G). This SNP is actually annotated as a splice acceptor variant, but that is based on annotation of an isoform that we did not detect in epithelial crypts. Instead, rs29595736 leads to an arginine (B6) to glycine (Balb) transition at amino acid 6 of full-length OCA-T1, which is just outside the POU2F3 binding site. Although we cannot rule out a change in protein function due to this SNP, its positioning just upstream of a transcriptional start site is more interesting from the perspective of isoform abundance. That said, 129S1/SvImJ mice, which phenocopy B6 mice, carry the Balb allele of rs29595736. Three other SNPs of interest are rs336266049, rs29736233, and rs37184010 (Fig. 6B, marked in red). These SNPs are all intronic, but they correlate with the tuft cell phenotypes of inbred strains; B6 and 129S1/SvImJ encode the same nucleotide, while Balb, FVB/NJ and C3H/HeJ all encode a different nucleotide. More work is needed to understand whether these and/or more distal SNPs impact isoform expression or tuft cell differentiation.

### Tuft cell abundance tunes sensitivity and kinetics of the tuft-ILC2 circuit

To understand the physiologic impact of low baseline tuft cell frequency and the role of *Pou2af2*, we used protists and helminths to activate the tuft-ILC2 circuit. While acute administration of succinate failed to activate the tuft-ILC2 circuit in Balb mice, *Tritrichomonas* protists chronically colonize mice from weaning and are perhaps more immunostimulatory than succinate alone. Nonetheless, adult Balb mice colonized with *Tritrichomonas* from birth failed to induce tuft cell hyperplasia. Responses in congenic Strain 3 mice were more variable, with only some mice developing hyperplasia despite elevated baseline tuft cell frequency in all mice. The lower expression of *Sucnr1* (Fig. 5B) and an unexplained ~60% lower protist burden (Fig. 7B) in Strain 3 mice likely kept them at or below the threshold for tuft-ILC2 circuit activation.

**Figure 7.**
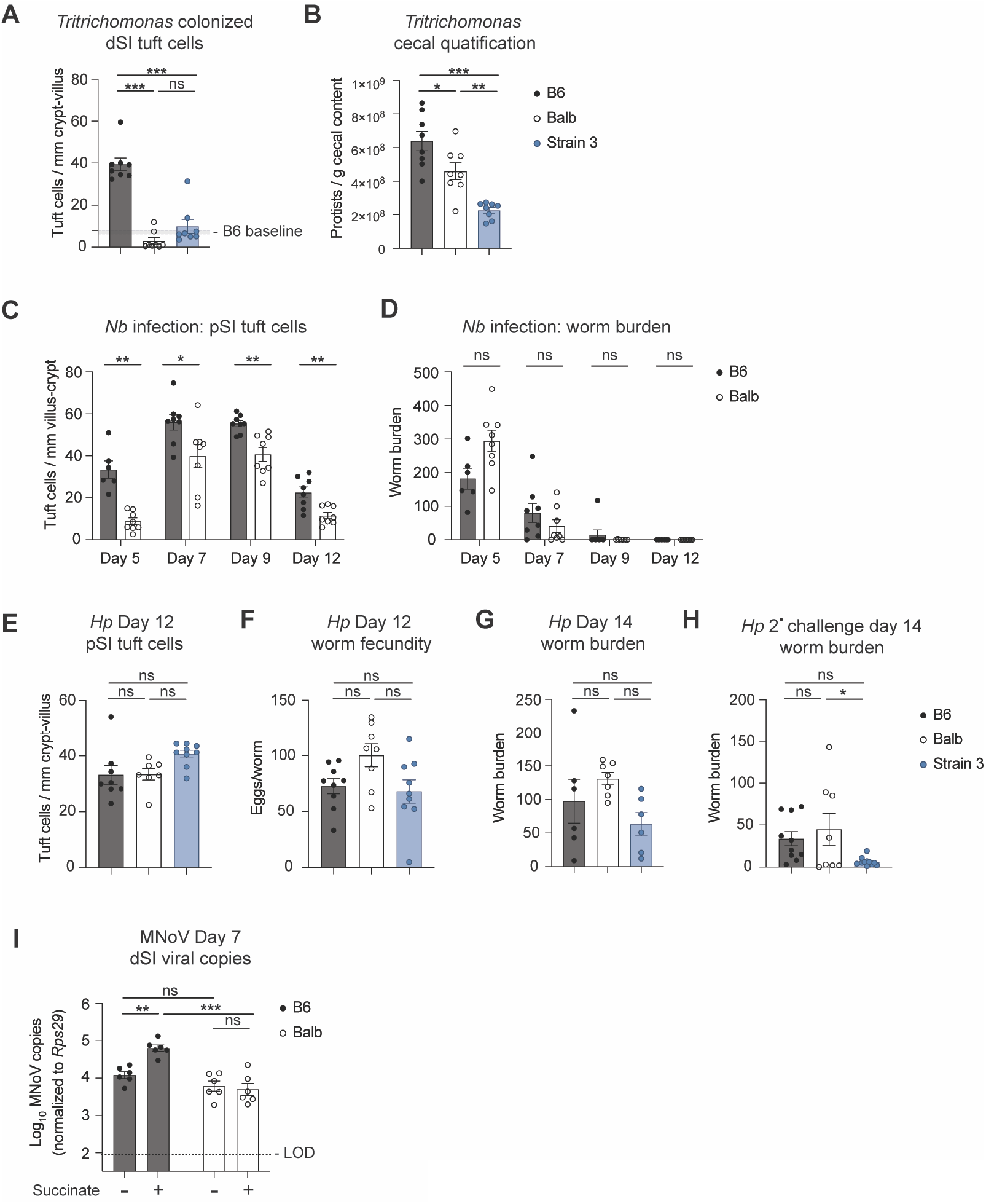
Tuft cell frequency at baseline tunes the kinetics and sensitivity of the tuft-ILC2 circuit. (**A**) Tuft cell quantification in the distal SI and (**B**) protist quantification in the cecal content of *Tritrichomonas* colonized mice. (**C**) Tuft cell quantification in the proximal SI and (**D**) worm burden in total intestine at the indicated time points post *Nb* infection. (**E**) Tuft cell quantification in the proximal SI on day 12 post *Hp* infection. (**F**) Overnight egg production by worms isolated from the proximal SI of mice 12 days post *Hp* infection. (**G**and **H**) Intestinal worm burden on day 14 of (G) primary or (H) secondary *Hp* infection 28 days after drug-cleared primary infection. (I) Mice were pretreated with 150mM sodium succinate or 300mM sodium chloride for 1 week prior to oral infection with murine norovirus (MNoV) CR6. Viral genome copies detected in the distal SI 7 days after CR6 infection. Dotted line represents limit of detection (LOD). In (A), shaded area indicates the 95% confidence interval of the mean for distal SI tuft cell quantification calculated from a large cohort of control B6 mice. In the graphs, each symbol represents an individual mouse from two or three pooled experiments. *p < 0.05, **p < 0.01, ***p < 0.001 by multiple t tests (C and D), by one-way ANOVA (A and B, E to H) or by two way ANOVA (I). n.s., not significant. Graphs depict mean +/− SEM. Also see Figure S7.

Next, we infected Balb and B6 mice with the helminth *Nippostrongylus brasiliensis* (*Nb*), an acute infection model that strongly activates the tuft-ILC2 circuit and is cleared within 7-8 days in B6 mice. Over the course of infection, Balb mice developed tuft cell hyperplasia, but with delayed kinetics compared to B6 (Fig 7C, fig. S7A). Balb mice had 50% higher worm burden on day 5 post infection, but complete worm clearance was not delayed (Fig. 7D). Therefore, although Balb mice start with fewer tuft cells, tuft-ILC2 circuit activation reaches a threshold required for *Nb* clearance and/or other mechanisms, such as a stronger adaptive Th2 responses, compensate for innate defects in Balb mice.

*Heligmosomoides polygyrus* (*Hp*) provides a model of long-term SI helminth infection, with clearance taking 6 weeks or more *(33)*. As mentioned previously, Balb mice clear *Hp* infection more rapidly than B6, likely due to a stronger adaptive type 2 immune response *(17, 34, 35)*, but the differences during early infection have not been well characterized. We wondered whether Strain 3 mice could benefit from both enhanced B6-like innate responses and a stronger Balb-like Th2 response. During primary infection, all three strains had equivalent tuft cell hyperplasia by day 12, with Strain 3 mice trending towards having more tuft cells compared to both B6 and Balb (Fig. 7E). However, worm fecundity (eggs laid per worm), worm burden, and total fecal egg counts trended lower in both B6 and Strain 3 mice, suggesting an earlier onset of protective immunity (Fig. 7F–G, fig. S7B). To test immune memory, we infected mice with *Hp* for 14 days, cleared infection with pyrantel pamoate, waited 28 days, and then challenged the mice with a secondary *Hp* infection. On Day 14 of challenge infection, Balb and Strain 3 mice had fewer worms in the intestine compared to B6 (Fig. 7F). Together these data demonstrate that innate tuft-ILC2 responses are delayed or even can develop and contribute to enhanced worm restriction and clearance. Congenic mice demonstrate both early B6-like and late Balb-like restriction.

In addition to sensing helminths and protists, intestinal tuft cells are the reservoir for murine norovirus strain CR6 and previous work has demonstrated that norovirus burden is regulated by type 2 signaling*(36)*. Accordingly, unmanipulated B6 mice had ~2-fold higher CR6 burdens than Balb. Treatment with succinate to mimic protist colonization increased ileal CR6 titers ~5-fold in B6 mice, but had no effect on titers in Balb mice (Fig. 7I). The two strains had similar norovirus titers in the colon regardless of succinate treatment (fig. S7D). In sum, baseline tuft cell frequency helps determine the sensitivity and kinetics of the innate tuft-ILC2 circuit. Balb mice maintain functional responses to helminth infection while ignoring *Tritrichomonas* colonization and lowering their norovirus burden.

## Discussion

Since the identification of the tuft-ILC2 circuit, numerous studies have uncovered ligands, receptors, and effector molecules that regulate this circuit. Much less progress has been made towards understanding cell intrinsic pathways by which epithelial stem cells commit to a tuft cell lineage, and how this process regulates tuft-ILC2 circuit activation. Here we identified differential *Pou2af2* isoform usage as a mechanism that establishes the baseline frequency of tuft cells in multiple tissues and tunes the sensitivity and kinetics of innate type 2 immunity in the SI.

We found that while unmanipulated Balb mice had fewer tuft cells at mucosal sites throughout the body, the B6 sequence from 50-67Mb on Chr9 was sufficient to restore tuft cell numbers to a B6 level in the SI and trachea (Fig. 4F–G, fig. S4D and F). Further, congenic mice carrying this interval develop hyperplasia when treated with succinate or, in some cases, when colonized with *Tritrichomonas* (Fig. 4H and Fig. 7A).

At 51.2 Mb on Chr9, adjacent to the QTL peak at 50.8 Mb, is *Pou2af2*, which was recently shown to encode a POU2F3 co-factor (OCA-T1), and to be necessary for tuft cell differentiation in the SI *(29)*. We found two isoforms of *Pou2af2* expressed in distal SI crypts, a full-length isoform and a shorter isoform, which lacks the POU2F3 interaction domain (Fig. 6A–B). Balb and Strain 1 SI crypts express significantly less of the functional full-length *Pou2af2* isoform compared to B6 or Strain 2-4 SI crypts (Fig. 6D), leading us to propose that *Pou2af2* isoform usage determines the number of tuft cells at baseline in the SI. What exactly determines isoform transcription is, as yet, unknown. There are several SNPs of interest within the *Pou2af2* locus, but transcription may be regulated by distal enhancers. The apparent co-regulation of *Pou2af2* and *Pou2af3* transcription in particular suggests a broader regulatory mechanism that may be revealed by analysis of 3D genome structure.

Interestingly, the entire region of mouse Chr9 from 26.7 to 54 Mb is syntenic with human Chr11, suggesting that the shared function and regulation of genes in this region is evolutionarily conserved. In addition to the *Pou2af* gene cluster, *Pou2f3* is located at 43Mb on mouse Chr9 and 120Mb on human Chr11. Intriguingly, *Pou2f3* has been linked to several human cancers, including small cell lung cancer and colon cancer*(37, 38)*. Particularly relevant to our findings, SNPs in or near *Pou2af2* have also been linked to colon cancer and tuft cell abundance through genome-wide association studies and *in silico* analysis*(39, 40)*. Further studies are needed to fully reveal the role of tuft cells and regulation of these genes in the context of human cancers and immunity. Other tuft cell signature genes in this syntenic region include *Nrgn* and *Dscaml1*, and further analysis may identify additional genes or regulatory elements that regulate tuft cell differentiation and function.

The identification of OCA-T1 (*Pou2af2*) and OCA-T2 (*Pou2af3*) as POU2F3 co-factors advanced our understanding of tuft cell differentiation*(29, 30)*. Here we add isoform usage as another layer of regulation. Together, these findings suggest interesting avenues for further study. In particular, how do OCA-T1, OCA-T2, and POU2F3 interact with each other and with other transcription factors thought to play a role in SI tuft cell differentiation, such as GFI1b, SOX4 and ATOH1*(41–44)*? In cell lines derived from human tuft-cell-like variants of small-cell lung cancer, deletion of *Pou2af2* results in decreased *Pou2f3* expression and vice versa, suggesting the two genes may impact each other’s expression*(29, 37)*. Whether this relationship is present in non-malignant tuft cells in human or mouse still needs to be elucidated. Lastly, our results from organoids, helminth infection, and succinate stimulation after rIL-25 priming suggest that unlike homeostatic differentiation, the response of Balb progenitors to IL-13 is similar to B6. How *Pou2af2* and *Pou2af3* are regulated in this context remains unknown.

*Pou2af2* isoform usage determines the baseline number of tuft cells, and this tunes the sensitivity and kinetics of the tuft-ILC2 circuit. During helminth infection, tuft cells secrete cysLTs, which potently activate ILC2s when paired with IL-25*(13)*. However, tuft cells do not produce cysLTs when stimulated with succinate*(13)*. Furthermore, the tuft cell deficit in Balb mice is more pronounced in the distal SI, where succinate sensing occurs, than in the proximal SI where helminths predominantly reside (Fig 1B). Ultimately, the integration of baseline tuft cell frequency and strength of signal sets the threshold for tuft-ILC2 circuit activation. Balb mice have few tuft cells but perhaps can overcome this defect with IL-25 and cysLT synergy downstream of helminth sensing. B6 mice likely rely on a higher baseline tuft cell frequency to sense the weaker, cysLT-independent, succinate signal. As a result, while adaptive immune responses in Balb mice are indeed skewed towards type 2 immunity, their innate type 2 immune response is attenuated, particularly in the distal SI. Rather than representing a defect, however, the lower baseline tuft cell frequency in Balb mice may be adaptive. Balb mice maintain functional responses to helminth pathogens while not expending the energy to remodel their epithelium in response to commensal protists. To date, no detrimental effects have been demonstrated in mice that fail to sense Tritrichomonads. Furthermore, the lack of tuft cell hyperplasia in Tritrichomonas-colonized Balb mice can lower their norovirus burden.

The link between tuft cells and immunity extends beyond helminth infection and protist colonization. In addition to expanding the niche for norovirus, acutely activating the tuft-ILC2 circuit results in worse outcomes for West Nile Virus infection in mice *(36, 45)*. Distal SI tuft cells can also sense bacterial-derived succinate, and in mice, giving succinate to increase tuft cell frequency reduces inflammation in models of ileitis*(11, 46)*. Perhaps relatedly, in Crohn’s disease, tuft cell frequency is lowest in areas of highest inflammation. Beyond the intestine, tracheal tuft cells sense bacterial ligands and regulate breathing, mucociliary clearance, and neuroinflammation *(47–49)*. In the gallbladder, tuft cells prevent inflammation, perhaps by inducing mucus production and smooth muscle contraction*(50, 51)*. Our data indicate that the Chr9 locus impacts baseline tuft cell frequency in multiple tissues, including the trachea. We expect that *Pou2af2* isoform usage, and associated tuft cell abundance, would influence tuft cell-mediated immune responses in these other contexts as well.

## Methods

### Experimental Animals

Mice aged 6 weeks and older were used for all experiments. Mice were age-matched within each experiment. Pooled results include both male and female mice of varying ages unless otherwise indicated. C57BL/6J (B6), and Balb/cJ (Balb) mice were bred in house or purchased from Jackson Laboratories. C3H/HeJ, FVB/NJ, 129S1/SvImJ and Swiss Webster mice were purchased from Jackson Laboratories. Congenic mice were generated and bred in house as described below. All mice were maintained in specific pathogen-free conditions at the University of Washington and were confirmed to be free of *Tritrichomonas* by microscopy and qPCR, unless specifically colonized for experimental purposes. All procedures were conducted within University of Washington IACUC guidelines under approved protocols.

### Quantitative trait locus mapping

We performed quantitative trait locus (QTL) mapping of succinate-induced tuft cell frequency using an F2 intercross between B6 and Balb. We generated Tn5-tagmented whole-genome sequencing libraries for 84 F2 hybrids and sequenced the samples to a depth of ~0.05x in a NextSeq 500/550 (75 cycles). Adapters were trimmed using Trimmomatic v0.36 (Bolger et al. 2014), and reads were aligned to the mm10 reference genome using BWA-MEM.

To impute genotypes, we generated a panel of SNPs between B6 and Balb using sequence variation data from the Mouse Genomes Project (Adams et al. 2015). SNPs that passed the following thresholds were included in the panel: MQ >= 60, DP between 40 and 140, GQ >= 60, and QUAL > 200. We genotyped each individual at all qualifying variant positions and conducted genotype imputation using Ancestry-HMM v0.94*(52)*. Genome-wide genotype probabilities from Ancestry-HMM were used to perform QTL analysis of succinate-induced tuft cell frequency using R/qtl. The code and results for this analysis are included as Data File S1.

### Congenic strain generation

We generated four congenic strains through six to eight generations of backcrossing to Balb to fine map the QTL on chromosome 9. Each generation, libraries were generated, sequenced, and aligned as described above. Genotypes were imputed using Ancestry-HMM v0.94. In each cohort, individuals were prioritized for continued backcrossing if recombination occurred within the congenic interval on chromosome 9. At a minimum, individuals chosen for breeding retained B6 ancestry in the chr9 locus and contained a high proportion of Balb ancestry outside the chr9 locus. Because we were unsuccessful at designing a method in which we could quantify succinate-driven tuft cell hyperplasia without euthanasia of the mouse, we selected breeders based only on their genotype and then phenotyped siblings with the same Chr9 genotypes for succinate responsiveness. After 6-8 generations of backcrossing, each congenic genome was homozygous for Balb DNA at all locations except the Chr9 locus, where they were homozygous for B6 DNA.

### Succinate Treatment

For succinate experiments mice were given 150mM or 250mM sodium succinate hexahydrate (Thermo) ad libitum in drinking water for the indicated amount of time.

### *In vivo* recombinant cytokine administration

IL-4 complexes were generated by incubating 2 μg mouse rIL-4 (R&D Systems) with 10 μg LEAF purified anti-mouse IL4 antibody (clone 11B11, Biolegend) per mouse for 30 min at room temperature. rIL-4 complex or 500ng rIL-25 were given for 3 consecutive days intraperitoneally in 200 ul PBS.

### Mouse Infection and Treatments

*H*. *polygyrus* and *N. brasiliensis* larvae were raised and maintained as previously described*(53, 54)*. Mice were infected by oral gavage with 200 *H*. *polygyrus* L3 or subcutaneously with 500 *N. brasiliensis* L3 and euthanized at the indicated time points to collect tissues for staining and/or to count worm burden. Worm burden was enumerated across the entire small intestine using a dissection microscope.

### *H. polygyrus* worm fecundity

Adapted from a previously described method*(55)*, 12 female worms were isolated from the proximal 5cm of the small intestine per mouse and individually cultured in 200ul plain RPMI 1640 with 200 U/mL penicillin and 200 μg/mL streptomycin in 96-well plates at 37°C. After 24 hours, eggs were counted, and eggs/worm were calculated.

### *H. polygyrus* fecal egg count

For fecal egg burdens, 2 to 3 fecal pellets were collected and weighed at time of euthanasia. Pellets were softened in PBS, homogenized with electric pestle, and transferred to 5mL H20 saturated with NaCl and eggs were counted using a McMaster’s Slide.

### Protist colonization

For protist colonization experiments, breeding pairs were colonized with *Tritrichomonas musculis* as previously described*(5)*. Pups from colonized breeding pairs were analyzed. Protist colonization was quantified by collecting and weighing cecal content at time of euthanasia, diluting in PBS and counting protists using a hemocytometer.

### Generation of murine norovirus stock

Stocks of murine norovirus (MNoV) strain CR6 were generated from molecular clones as previously describe*(56)* except for a modified virus concentration protocol. Briefly, plasmids encoding the viral genomes were transfected into 293T cells to generate infectious virus, which was subsequently passaged on BV2 cells. After two passages, BV2 cultures were frozen and thawed to liberate virions. Virus was concentrated by centrifugation in a 100,000 MWCO ultrafiltration unit (Vivaspin, Sartorius). Titers of virus stocks were determined by plaque assay on BV2 cells*(57)*.

### MNoV infections, sample collection and quantification

Mice received either 150mM sodium succinate or 300mM sodium chloride in the drinking water for 7 days prior to infection with CR6 and continued to receive treatment water until time of harvest. 6-week-old mice were orally inoculated with 10^6^ PFU of CR6 in a volume of 25μl. 7 days post Cr6 infection tissues and stool were harvested into 2-ml tubes (Sarstedt) with 1-mm-diameter zirconia/silica beads (Biospec). Samples were frozen and stored at −80C until RNA extraction. As previously described*(58)*, RNA from tissues was isolated using TRI Reagent with a Direct-zol-96 RNA kit (Zymo Research) according to the manufacturer’s protocol. 5μl of RNA was used for cDNA synthesis with the ImPromII reverse transcriptase system (Promega). MNoV TaqMan assays were performed, using a standard curve for determination of absolute viral genome copies, as described previously *(59)*. qPCR for housekeeping gene *Rps29* was performed as previously described *(60)*. All samples were analyzed with technical duplicates.

### Intestinal tissue fixation and staining

Intestinal tissues were flushed with PBS and fixed in 4% paraformaldehyde for 4 hours at 4°C. Tissues were washed with PBS and incubated in 30% (w/v) sucrose overnight at 4°C. Samples were then coiled into “Swiss rolls” and embedded in Optimal Cutting Temperature Compound (Tissue-Tek) and sectioned at 8 μm on a CM1950 cryostat (Leica). Immunofluorescent staining was performed in PBS with 1% BSA at room temperature (RT) as follows: 1 h 5% goat serum, 1 h primary antibody (αDCLK1, Abcam ab31704), 40 min goat anti-rabbit IgG F(ab’)2-AF594 secondary antibody (Invitrogen) and mounted with Vectashield plus DAPI (Vector Laboratories). Images were acquired with an Axio Observer A1 (Zeiss) microscope with a 10X A Plan objective. Tuft cell frequency was calculated using ImageJ software to manually quantify DCLK1^+^ cells per millimeter of crypt-villus axis. Four 10x images of the Swiss roll were analyzed for each replicate and at least 25 total villi were counted.

### Tracheal tuft cell staining and quantification

Tracheas were harvested and connective tissue was removed. Tracheas were opened longitudinally and washed 5 times in 5% FBS/10mM DTT/0.05% Tween-20/HBSS, vortexing for 5 seconds, to remove mucus. Tracheas were stretched out by pinning to SylGard-coated well of 6 well plate and fixed for 1 hr on ice in Cytofix/Cytoperm buffer (BD Biosciences). Immunofluorescent staining was performed in PBS with 0.25% Triton X-100 at 4°C as follows: 24 h 10% goat serum, 24 to 36 h primary antibody (αDCLK1, Abcam ab31704), 2 h goat anti-rabbit IgG F(ab’)2-AF488 secondary antibody (Invitrogen), 15 min DAPI (1:1000), and mounted with Vectashield (Vector Laboratories). Images were acquired with a Nikon eclipse Ti microscope using a CSU-W1 spinning disc confocal with a Plan Apo λ 20X objective. 5 images were collected per sample and tuft cells were quantified using QuPath cell detection software.

### Intestinal single-cell tissue preparation

For single cell epithelial preparations from small intestines or cecum, tissues were flushed with PBS, Peyer’s patches removed, opened longitudinally, and rinsed with PBS to remove intestinal contents and mucus. Intestinal tissue was cut into 2-5 cm pieces and cecum was cut into 5-6 strips. Tissues were incubated rocking at 37°C for 10 min in 10ml HBSS (Ca^+2^/Mg^+2^-free) supplemented with 3mM EDTA and 1mM HEPES. Tissues were vortexed thoroughly and released epithelial cells were passed through a 70 μm filter. Tissues were then incubated in fresh EDTA/HBSS solution and incubation, vortexing and filtering was repeated for a total of 3 rounds. Supernatants were pooled and washed once before staining for flow cytometry.

For lamina propria preparations, small intestinal tissue was processed as above to remove the epithelial fraction. Tissues were then incubated in 10ml RPMI 1640 supplemented with 20% FCS, 1mM HEPES, 0.05 mg/ml DNase I (Sigma Aldrich), and 1 mg/mL Collagenase A (Sigma Aldrich), shaking at 37°C for 30 minutes. Tissues were vortexed and cells were passed through a 100 μm filter, then a 40 μm filter. Cells were then washed and stained for flow cytometry.

### Thymus single-cell tissue preparation

For thymus epithelial preparations, protocol was adapted from previously described procedure*(2)*. Briefly, thymi cleaned of fat were minced with a razor blade. Tissue was incubated in 37°C water bath for 12 min in 4 ml of digestion medium containing 2% FBS, 100 μ/ml DNase I (Sigma Aldrich) and 100 μ/ml liberase TM (Sigma Aldrich) in DMEM. At 12 min, tubes were spun briefly to pellet undigested fragments and the supernatant was moved to 20 ml of 0.5% BSA, 2 mM EDTA in PBS on ice. The DNAse/Liberase digestion was repeated twice for a total of three 12-min digestion cycles. The single-cell suspension was pooled, pelleted and resuspended in 50% Percoll (Sigma Aldrich), underlaid with 90% Percoll, and centrifuged at 2,000 rpm for 15 min at 20°C. The 50/90 interphase of the Percoll gradient was collected, washed, and stained for flow cytometry as described below.

### Organoid Culture

Small intestinal crypt-derived organoids were grown as described with modifications described below*(61)*. Briefly, distal small intestine was isolated and villi manually scraped off with a glass coverslip. Tissue was then washed three times in cold PBS with vigorous shaking before 30 minute 4 °C incubation in 2mM EDTA to release epithelial crypts, which were washed in PBS and filtered through a 70 μm strainer. Pelleted crypts were resuspended in Matrigel and plated at 400-500 crypts per well in a prewarmed plate, incubated at 37°C for 5 minutes to allow for Matrigel solidification, and complete organoid media added. Organoid media was composed of DMEM/F12 supplemented with 2mM glutamine, 100 U/mL penicillin, 100mg/mL streptomycin, 100ug/mL Normacin (InvivoGen), 10mM HEPES, 1X N2 supplement (Life Technologies), 1X B27 supplement (Life Technologies), 500mM N-acetylcysteine, 50μg/ml mEGF, and replacing recombinant R-spondin with supernatants from R-spondin expressing L-cells and replacing recombinant Noggin with supernatants from Noggin expressing cells. Crypts were harvested from distal (last 10cm) small intestine of naive mice and plated on day 0. On day 3 and day 5, media was replaced. Organoids were treated with 2.5 ng/ml recombinant IL-13 on day 1, 3 and 5. On day 7 organoids were harvested for passage or analysis. Organoids were passaged by washing in room temperature PBS to remove Matrigel. Next, organoids were sheared with a 28G insulin syringe, washed and resuspended in fresh Matrigel. Generally, organoids were passaged at 1 well to 3-5 well ratio depending on number of organoids present.

For flow cytometry, organoids were resuspended in 1X TrypLE (Gibco). Organoids were sheared with a 28G insulin syringe, incubated for 10min at room temperature, cells washed, and then stained for flow cytometry as described below. Tuft cells were identified as CD45^−^ EpCAM^+^ CD24^+^ DCLK1^+^. For qPCR, organoids were incubated in Cell Recovery Solution (Corning) for 30 min at 4°C to remove Matrigel. Organoids were washed 2 times with PBS, pelleted and resuspended in RLT Plus buffer. RNA was isolated using the Mini Plus RNeasy kit (Qiagen) following manufacture’s protocol.

### Flow cytometry and cell sorting

Single cell suspensions from tissues or organoids were prepared as described above. For flow cytometry, samples were stained with Zombie Violet (BioLegend) in PBS for live/dead exclusion and stained in PBS + 3% FBS with antibodies to surface markers. Next, cells were fixed and permeabilized using the eBioscience™ Foxp3 / Transcription Factor Staining Buffer Set, following manufacturer’s instructions for staining either cytosolic proteins (DCLK1) or nuclear proteins (GATA3 and Ki67). When cell counts were needed, counting beads (Invitrogen) were added prior to running flow cytometry. Samples were run on a Canto RUO or Symphony A3 (BD Biosciences) and analyzed with FlowJo 10 (Tree Star). Samples were FSC-A/SSC-A gated to exclude debris, FSC-A/FSC-H gated to select single cells and gated to exclude dead cells.

For cell sorting of ILC2s or tuft cells, single cell suspensions were prepared as described above. Cells were stained in PBS + 3% FBS with antibodies to surface markers and stained with DAPI for live/dead exclusion. Samples were sorted on an Aria II (BD Biosciences). Samples were FSC-A/SSC-A gated to exclude debris, FSC-A/FSC-H gated to select single cells and gated to exclude dead cells.

### ILC2 Stimulation Assay

Small intestinal lamina propria ILC2s were isolated from mice and sorted as described. Sorted cells were plated at 5000 cells per well in a 96 well plate and incubated at 37°C overnight in 10 ng/ml IL-7 (R&D Systems) and basal media composed of high glucose DMEM supplemented with non-essential amino acids, 10% FBS, 100 U/mL penicillin, 100mg/mL streptomycin, 10mM HEPES, 1mM sodium pyruvate, 100μM 2-mercaptoethanol, and 2mM L-glutamine. The next morning, media was replaced with fresh media and 10 ng/ml IL-7, and cells were stimulated with the indicated agonist. After a six-hour stimulation at 37°C, supernatant was collected and analyzed by cytokine bead array as described below. Cells were resuspended in fresh basal media with 10ng/ml IL-7 and incubated for an additional 48 hrs. Cells were washed, stained for intracellular Ki67 as described above. Cytokine levels in supernatants collected from cultured ILC2s were measured using Enhanced Sensitivity Flex Sets (BD Biosciences) for mouse IL-5 and IL-13 according to the manufacturer’s protocol. Data was collected on a LSR II (BD Biosciences).

### Quantitative RT-PCR

Crypts from distal small intestine were isolated as described in the organoid culture methods. After filtering crypt suspension with 70 um filter, crypts were washed in PBS two times, pelleted and resuspended in RLT Buffer. RNA was isolated using the Mini Plus RNeasy kit (Qiagen) according to manufacturer’s instructions and reverse transcribed using SuperScript II (Thermo) following manufactures’ protocol. cDNA was used as template for quantitative PCR with PowerUP SYBR Green (Thermo) on a Via7 cycler (Applied Biosystems). Transcripts were normalized to *Rps17* (40S ribosomal protein S17) expression. Primer sequences listed in Table S2.

### RNA Sequencing and Analysis

150 tuft cells were sorted as CD45^lo^ EpCAM^+^SigF^+^CD24^+^ directly into lysis buffer from the SMART-Seq v4 Ultra Low Input RNA Kit (Takara) and cDNA was generated following manufacturer’s instructions. Four biological replicates were collected for each genotype. Each biological replicate represents one mouse. Next-generation sequencing was performed by the Benaroya Research Institute Genomics Core. Sequencing libraries were generated using the Nextera XT library preparation kit with multiplexing primers, according to manufacturer’s protocol (Illumina), and library quality was assessed using the Tapestation (Agilent). High throughput sequencing was performed on NextSeq 2000 (Illumina), sequencing dual-indexed and paired-end 59 base pair reads. All samples were in the same run with target depth of 5 million reads to reach adequate depth of coverage.

Processing and analysis of the raw sequencing reads was performed using the DIY.Transcriptomics (diytranscriptomics.com) pipeline, with experiment-specific modifications. Raw reads were mapped to the mouse reference transcriptome using Kallisto, version 0.46.2. The quality of raw reads, as well as the results of Kallisto mapping were analyzed using fastqc and multiqc. Kallisto outputs were read into an r environment and annotated using Biomart. Samples were filtered to exclude genes with counts per million = 0 in 4 or more samples and genes annotated as pseudogenes. Finally, samples were normalized to each other. To identify differentially expressed genes, precision weights were first applied to each gene based on its mean-variance relationship using VOOM, then data was normalized using the TMM method in EdgeR. Linear modeling and bayesian stats were employed via Limma to find genes that were up-or down-regulated by 2-fold or more, with a false-discovery rate (FDR) of 0.05. The code and results for this analysis are included as Data File S2.

### 5’ rapid amplification of cDNA ends

RNA isolated from distal SI crypts was obtained as described above. For 5’ rapid amplification of cDNA ends assay, the SMARTer RACE 5’/3’ Kit (Takara) was used following manufacturer’s protocol and using the following primer: *5*’-GATTACGCCAAGCTTGGTGGGCGGTAGTCTCCATAGGGCTCAGC-*3*’.

### Quantification and Statistical Analysis

All experiments were performed using randomly assigned mice without investigator blinding. All data points and “n” values reflect biological replicates (i.e. mice). No data were excluded. Statistical analysis was performed as noted in figure legends using Prism 7 (GraphPad) software. Graphs show mean +/− SEM.

## Acknowledgements

We thank all members of the von Moltke lab for helpful discussion and input on this manuscript. We thank D. Hailey and the Garvey Cell Imaging Lab in the Institute for Stem Cell & Regenerative Medicine for microscopy support; the mouse husbandry staff in the UW SLU vivarium; V. Gersuk, K. O’Brien and the Benaroya Research Institute Genomics Core for help with RNA sequencing; and M. F. Fontana for helpful comments on the manuscript. Flow cytometry data was acquired through the University of Washington, Cell Analysis Facility Shared Resource Lab, with NIH award 1S10OD024979-01A1 funding for the Symphony A3. M.S.N. was supported by a University of Washington Immunology Training Grant T32 AI106677 and by the UW Immunology Department Titus Fellowship. Work at Columbia University was supported by NIH R35GM143051, Sloan Foundation Fellowship, Klingenstein-Simons Fellowship (all to A.B.). A.B. and J.v.M. are Searle Scholars. J.v.M. is a Burroughs Wellcome Investigator in the Pathogenesis of Infectious Disease. Work at Washington University was supported by R01AI139314 (MTB) and F31AI167499 (EAK). Work at the University of Washington was supported by NIH DP2OD024087 and R01AI167923.

## Author contributions

MSN designed and performed experiments, analyzed data, and wrote the paper with JVM. LMW and DLJ assisted with experiments at the University of Washington. NN performed Tn5-tagmented whole-genome sequencing, genotype imputation for congenic strain generation and assisted with QTL analysis, with supervision by AB. EAK performed murine norovirus experiments and data analysis with supervision and funding provided by MTB. AB acquired funding and provided resources for Tn5-sequencing. JVM conceived of and supervised the study, performed experiments, analyzed data, and wrote the paper with MSN.

## Figure legends

**Supplemental Figure 1.**
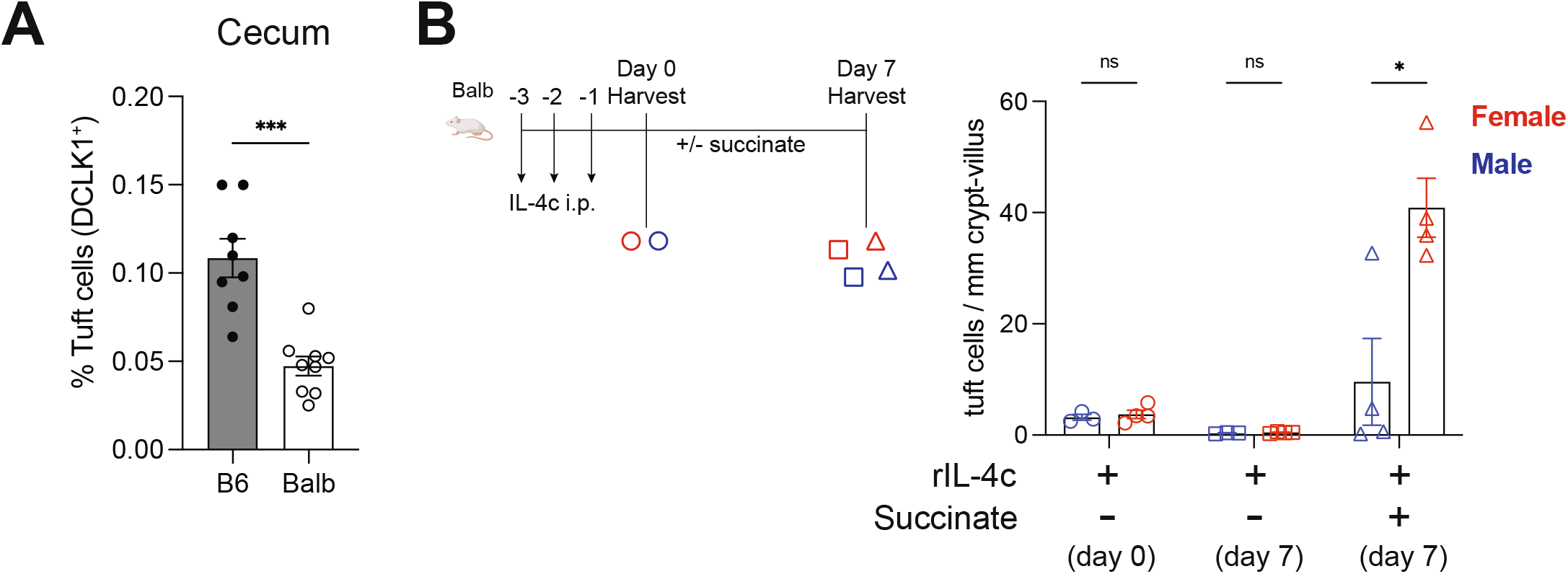
Balb mice have fewer tuft cells at baseline and rIL-4c priming leads to sex specific activation of the tuft-ILC2 circuit. (**A**) Quantification of cecal tuft cells by flow cytometry. (**B**) Data from Figure 1G separated by sex. In the graphs, each symbol represents an individual mouse from two or three pooled experiments. *p < 0.05, **p < 0.01, ***p < 0.001 by Mann-Whitney (A) or multiple t tests (B). n.s., not significant. Graphs depict mean +/− SEM.

**Supplemental Figure 2.**
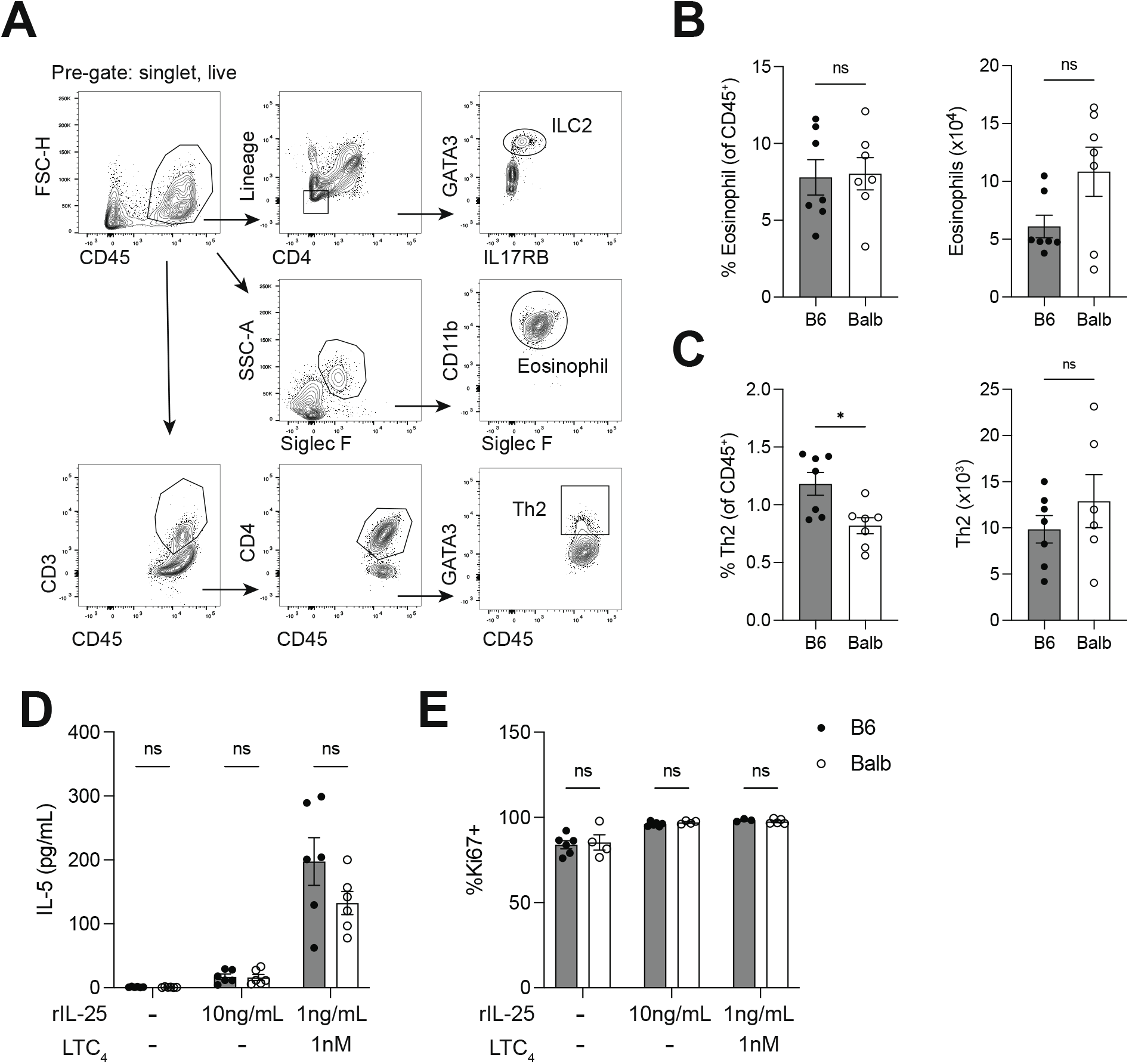
Equivalent numbers and responses of small intestinal type 2 immune cells in Balb and B6 mice. (**A**) Gating strategy for identification of ILC2s, eosinophils and GATA3^+^ Th2s from SI lamina propria of naive mice. (**B** and **C**) Percentage and absolute number of (B) eosinophils and (C) Th2 cells. (**D**) IL-5 concentration in the supernatant following 6-h *in vitro* culture of SI ILC2s with the indicated concentrations of rIL-25 and LTC_4_ and (**E**) Ki67 expression 2 days after stimulation. In the graphs, each symbol represents an individual mouse from two pooled experiments. *p < 0.05, **p < 0.01, ***p < 0.001 by Mann-Whitney (B and C) or by multiple t tests (D and E). n.s., not significant. Graphs depict mean +/− SEM.

**Supplemental Figure 3.**
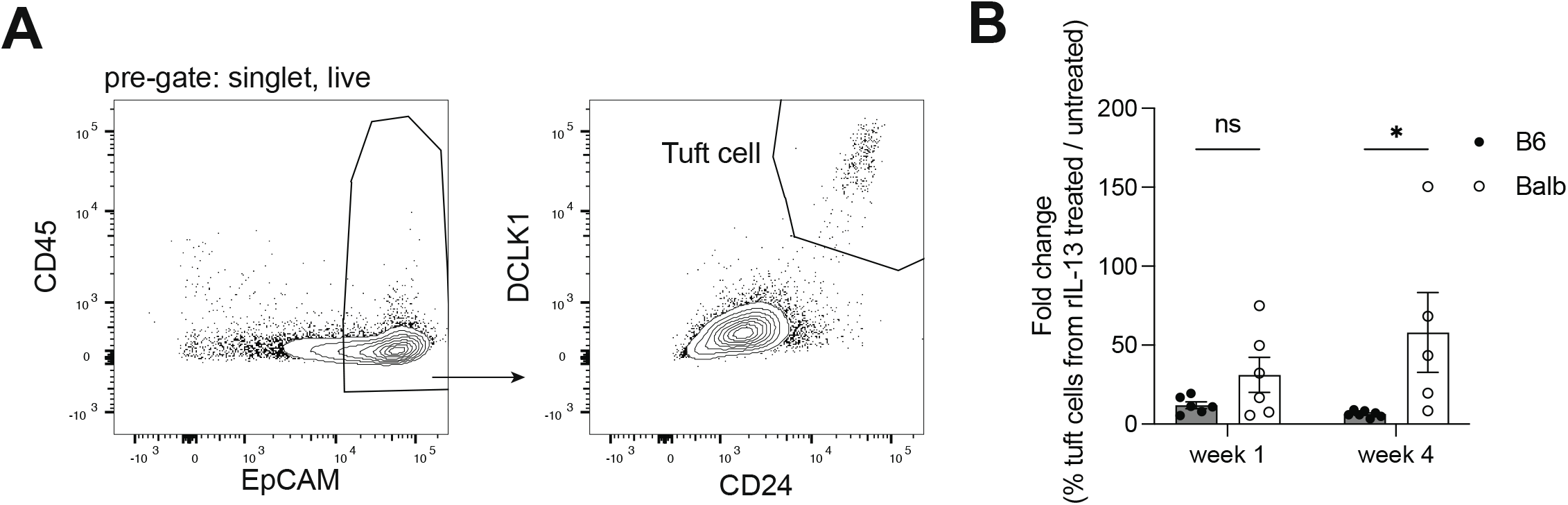
Organoid analysis. (**A**) Gating strategy for identification of tuft cells from SI organoids. (**B**) Fold change in % tuft cells from rIL-13 treated over untreated organoids derived from the same biological replicate. In the graphs, each symbol represents a biological replicate based on the average of 2 to 3 technical replicates, from three to six pooled experiments. *p < 0.05, **p < 0.01, ***p < 0.001 by multiple t tests (B). n.s., not significant. Graphs depict mean +/− SEM.

**Supplemental Figure 4.**
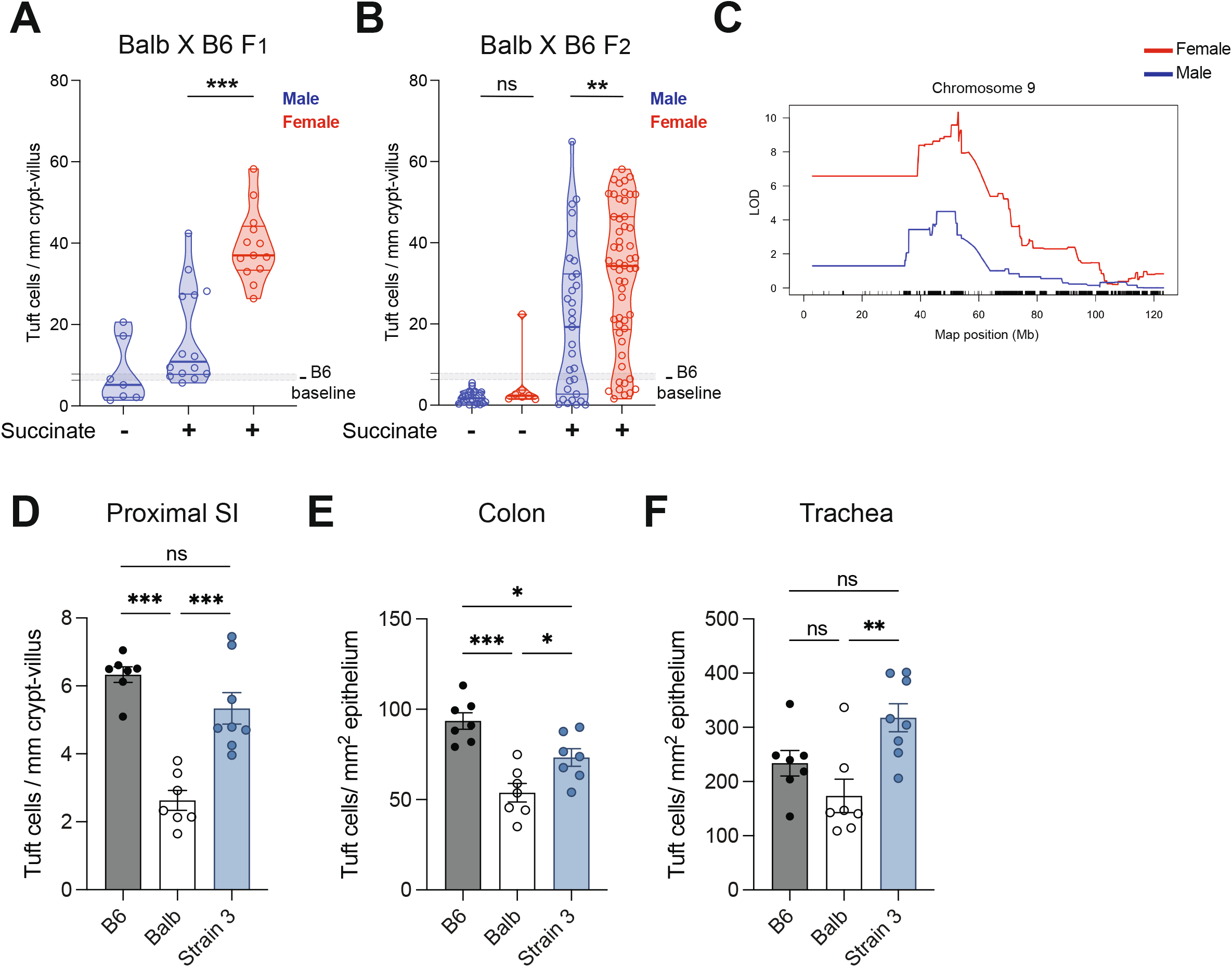
Sex effect in Balb x B6 F1 and F2 mice and recovery of tuft cell abundance in Strain 3 congenic. (**A** and **B**) Tuft cell quantification in dSI of Balb X B6 (A) F1 and (B) F2 mice by sex. (**C**) Chromosome 9 QTL mapping of succinate induced tuft cell hyperplasia in Balb X B6 F2 by sex. (**D**, **E** and **F**) Tuft cell quantification in the (D) proximal SI, (E) colon and (F) trachea by immunofluorescence. B6 and Balb data in (F) are also represented in Figure 1B. In the graphs, each symbol represents an individual mouse from three or more pooled experiments. *p < 0.05, **p < 0.01, ***p < 0.001 by multiple t tests (A and B) and by one-way ANOVA (D, E and F). n.s., not significant. Graphs depict mean +/− SEM.

**Supplemental Figure 5.**
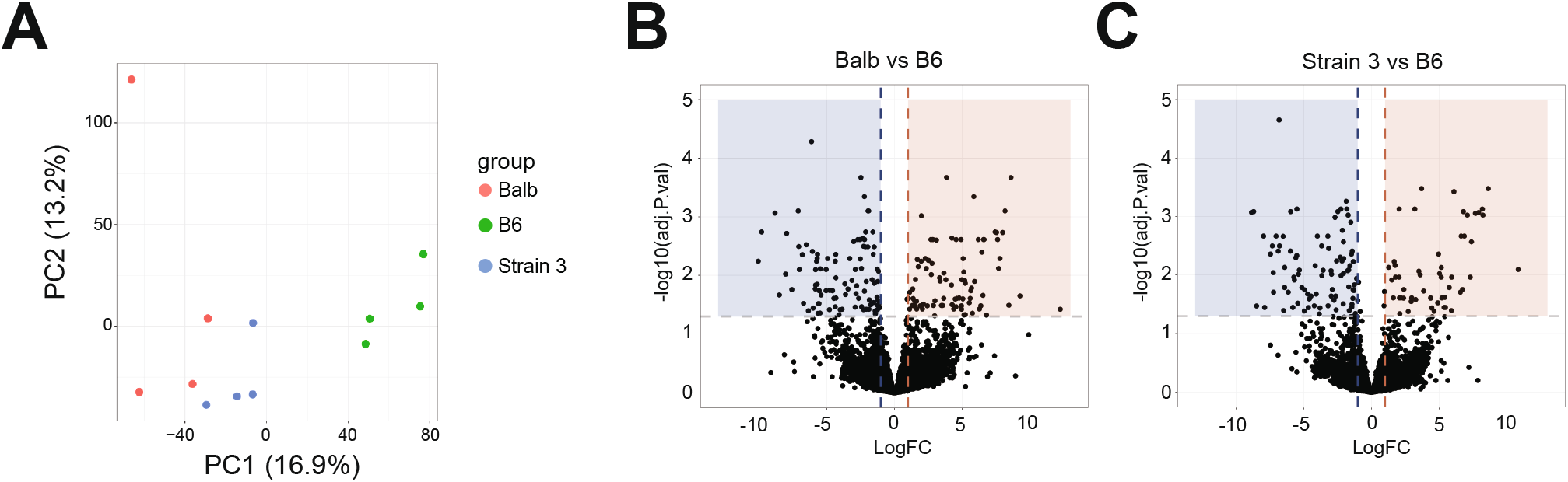
mRNA sequencing of mature tuft cells from B6, Balb and Strain 3 mice. (**A**) Unsupervised PCA of gene expression. (**B** and **C**) Volcano plots of (B) Balb vs B6 and (C) Congenic vs B6. The samples in this figure were all analyzed in one sequencing run.

**Supplemental Figure 6.**
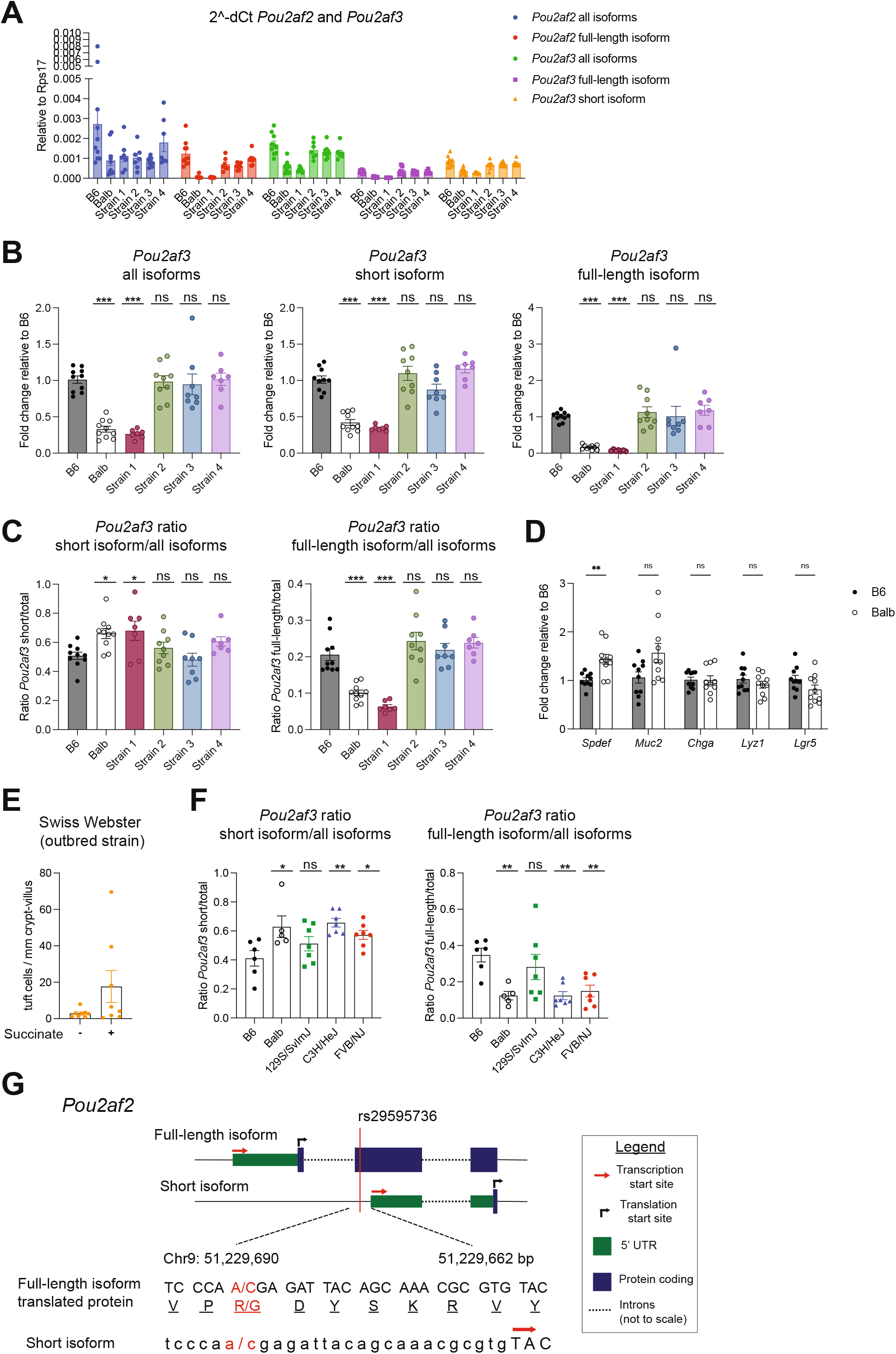
*Pou2af3* isoform expression follows similar pattern as *Pou2af2*. (**A**) Real-time PCR quantification of indicated genes/isoforms normalized to *Rps17* (housekeeping gene) from distal SI crypts. (**B**) *Pou2af3* isoform expression normalized to B6 and (**C**) *Pou2af3* isoform ratios. (**D**) Real-time PCR quantification of indicated genes normalized to B6. (**E**) Tuft cell quantification in dSI of Swiss Webster mice. (**F**) *Pou2af3* isoform ratios from indicated strains. (**G**) Depiction of SNP rs29595736 in *Pou2af2* isoforms and translated protein. In the graphs, each symbol represents an individual mouse from two or three pooled experiments. *p < 0.05, **p < 0.01, ***p < 0.001 by one-way ANOVA (B, C and F) with comparison to B6 and by multiple t tests (D). Graphs depict mean +/− SEM.

**Supplemental Figure 7.**
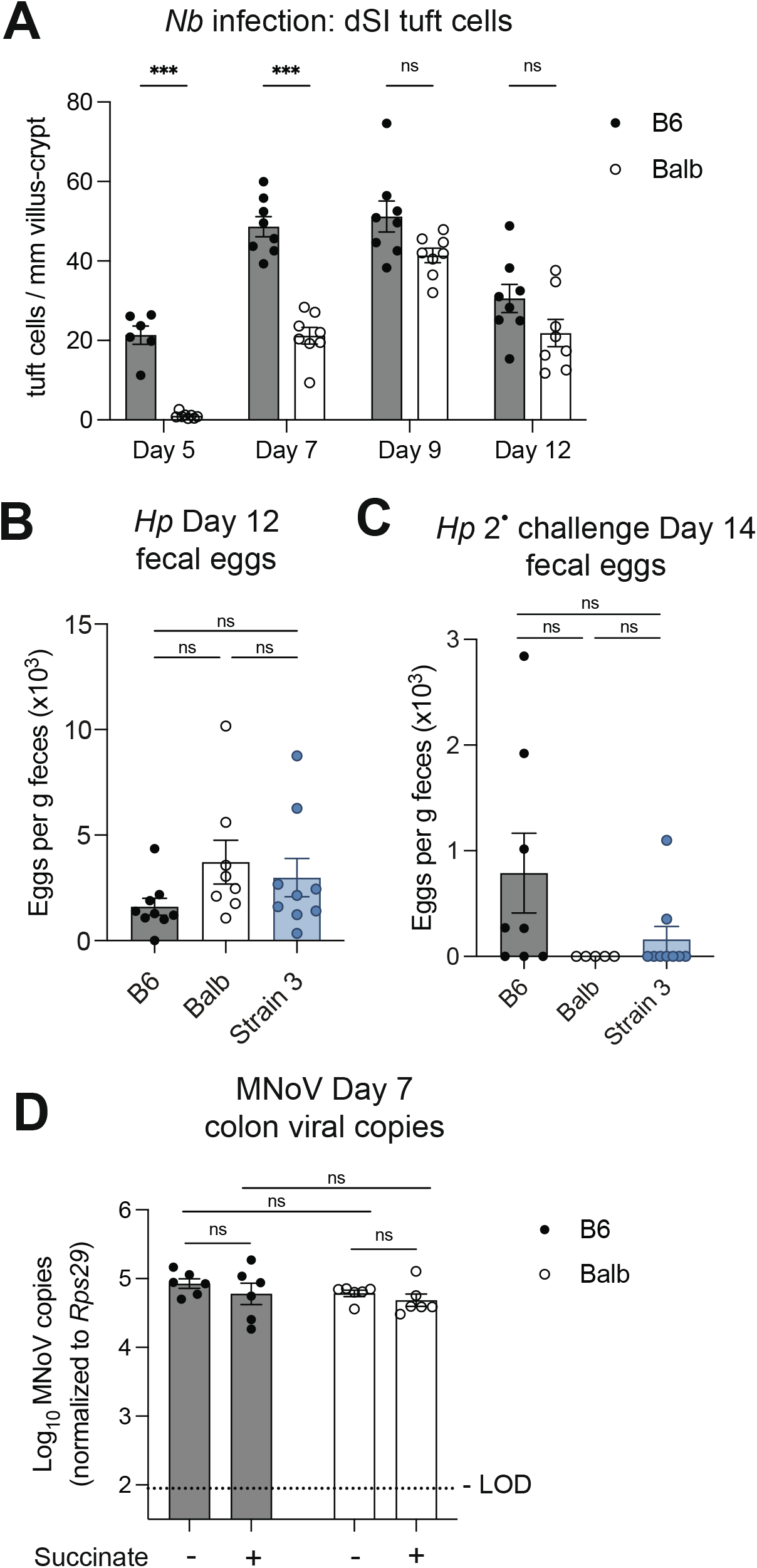
Tuft cell frequency at baseline tunes the kinetics and sensitivity of the tuft-ILC2 circuit. (**A**)Tuft cell quantification in the distal SI at the indicated time points post *Nb* infection. (**B and C**) Eggs per gram feces quantified from mice (B) 12 days post primary *Hp* infection or (C) 14 days post challenge *Hp* infection. (**D**) Mice were pretreated with 150mM sodium succinate or 300mM sodium chloride for 1 week prior to oral infection with murine norovirus (MNoV) CR6. Viral genome copies detected in the colon 7 days after CR6 infection. Dotted line represents limit of detection (LOD). In the graphs, each symbol represents an individual mouse from two pooled experiments. *p < 0.05, **p < 0.01, ***p < 0.001 by multiple t tests (A), by one-way ANOVA (B-C) and by two-way ANOVA (D). n.s., not significant. Graphs depict mean +/− SEM.

## References

1. C. E. O’Leary, C. Schneider, R. M. Locksley, Tuft Cells-Systemically Dispersed Sensory Epithelia Integrating Immune and Neural Circuitry. Annu. Rev. Immunol. 37, 47–72 (2019).

2. C. N. Miller, I. Proekt, J. von Moltke, K. L. Wells, A. R. Rajpurkar, H. Wang, K. Rattay, I. S. Khan, T. C. Metzger, J. L. Pollack, A. C. Fries, W. W. Lwin, E. J. Wigton, A. V. Parent, B. Kyewski, D. J. Erle, K. A. Hogquist, L. M. Steinmetz, R. M. Locksley, M. S. Anderson, Thymic tuft cells promote an IL-4-enriched medulla and shape thymocyte development. Nature 559, 627–631 (2018).

3. C. Bornstein, S. Nevo, A. Giladi, N. Kadouri, M. Pouzolles, F. Gerbe, E. David, A. Machado, A. Chuprin, B. Tóth, O. Goldberg, S. Itzkovitz, N. Taylor, P. Jay, V. S. Zimmermann, J. Abramson, I. Amit, Single-cell mapping of the thymic stroma identifies 25-producing tuft epithelial cells. Nat. 2018 5597715 559, 622–626 (2018).

4. C. Bezençon, A. Fürholz, F. Raymond, R. Mansourian, S. Métairon, J. Le Coutre, S. Damak, Murine intestinal cells expressing Trpm5 are mostly brush cells and express markers of neuronal and inflammatory cells. J. Comp. Neurol. 509, 514–525 (2008).

5. M. S. Nadjsombati, J. W. McGinty, M. R. Lyons-Cohen, J. B. Jaffe, L. DiPeso, C. Schneider, C. N. Miller, J. L. Pollack, G. A. Nagana Gowda, M. F. Fontana, D. J. Erle, M. S. Anderson, R. M. Locksley, D. Raftery, J. von Moltke, Detection of Succinate by Intestinal Tuft Cells Triggers a Type 2 Innate Immune Circuit. Immunity 49, 33–41.e7 (2018).

6. N. Barker, Adult intestinal stem cells: Critical drivers of epithelial homeostasis and regeneration Nat. Rev. Mol. Cell Biol. 15, 19–33 (2014).

7. F. Gerbe, E. Sidot, D. J. Smyth, M. Ohmoto, I. Matsumoto, V. Dardalhon, P. Cesses, L. Garnier, M. Pouzolles, B. Brulin, M. Bruschi, Y. Harcus, V. S. Zimmermann, N. Taylor, R. M. Maizels, P. Jay, Intestinal epithelial tuft cells initiate type 2 mucosal immunity to helminth parasites. Nature 529, 226–230 (2016).

8. J. von Moltke, M. Ji, H.-E. Liang, R. M. Locksley, Tuft-cell-derived IL-25 regulates an intestinal ILC2–epithelial response circuit. Nature 529, 221–225 (2016).

9. M. R. Howitt, S. Lavoie, M. Michaud, A. M. Blum, S. V Tran, J. V Weinstock, C. A. Gallini, K. Redding, R. F. Margolskee, L. C. Osborne, D. Artis, W. S. Garrett, Tuft cells, taste-chemosensory cells, orchestrate parasite type 2 immunity in the gut. Science (80-.). 351, 1329–1333 (2016).

10. C. Schubart, B. Krljanac, M. Otte, C. Symowski, E. Martini, C. Günther, C. Becker, C. Daniel, D. Voehringer, Selective expression of constitutively activated STAT6 in intestinal epithelial cells promotes differentiation of secretory cells and protection against helminths. Mucosal Immunol. 2018 122 12, 413–424 (2018).

11. W. Lei, W. Ren, M. Ohmoto, J. F. Urban, I. Matsumoto, R. F. Margolskee, P. Jiang, Activation of intestinal tuft cell-expressed Sucnr1 triggers type 2 immunity in the mouse small intestine. Proc. Natl. Acad. Sci. U. S. A. 115, 5552–5557 (2018).

12. C. Schneider, C. E. O’Leary, J. von Moltke, H.-E. Liang, Q. Y. Ang, P. J. Turnbaugh, S. Radhakrishnan, M. Pellizzon, A. Ma, R. M. Locksley, A Metabolite-Triggered Tuft Cell-ILC2 Circuit Drives Small Intestinal Remodeling. Cell 174, 271–284.e14 (2018).

13. J. W. McGinty, H. A. Ting, T. E. Billipp, M. S. Nadjsombati, D. M. Khan, N. A. Barrett, H. E. Liang, I. Matsumoto, J. von Moltke, Tuft-Cell-Derived Leukotrienes Drive Rapid Anti-helminth Immunity in the Small Intestine but Are Dispensable for Anti-protist Immunity. Immunity 52, 528–541.e7 (2020).

14. F. P. Heinzel, M. D. Sadick, B. J. Holaday, R. L. Coffman, R. M. Locksley, Reciprocal expression of interferon γ or interleukin 4 during the resolution or progression of murine leishmaniasis. Evidence for expansion of distinct helper T cell subsets. J. Exp. Med. 169, 59–72 (1989).

15. R. M. Locksley, P. Scott, Helper T-cell subsets in mouse leishmaniasis: induction, expansion and effector function. Immunol. Today 12, A58–A61 (1991).

16. W. Hartmann, B. Blankenhaus, M. L. Brunn, J. Meiners, M. Breloer, Elucidating different pattern of immunoregulation in BALB/c and C57BL/6 mice and their F1 progeny. Sci. Rep. 11, 1–14 (2021).

17. K. J. Filbey, J. R. Grainger, K. A. Smith, L. Boon, N. Van Rooijen, Y. Harcus, S. Jenkins, J. P. Hewitson, R. M. Maizels, Innate and adaptive type 2 immune cell responses in genetically controlled resistance to intestinal helminth infection. Immunol. Cell Biol. 92, 436–448 (2014).

18. L. A. Reynolds, K. A. Smith, K. J. Filbey, Y. Harcus, J. P. Hewitson, S. A. Redpath, Y. Valdez, M. J. Yebra, B. Brett Finlay, R. M. Maizels, Commensal-pathogen interactions in the intestinal tract lactobacilli promote infection with, and are promoted by, helminth parasites. Gut Microbes 5, 522–532 (2014).

19. J. M. Behnke, F. A. Iraqi, J. M. Mugambi, S. Clifford, S. Nagda, D. Wakelin, S. J. Kemp, R. L. Baker, J. P. Gibson, High resolution mapping of chromosomal regions controlling resistance to gastrointestinal nematode infections in an advanced intercross line of mice. Mamm. Genome 17, 584–597 (2006).

20. J. D. Gorham, M. L. Güler, R. G. Steen, A. J. Mackey, M. J. Daly, K. Frederick, W. F. Dietrich, K. M. Murphy, Genetic mapping of a marine locus controlling development of T helper 1/T helper 2 type responses. Proc. Natl. Acad. Sci. U. S. A. 93, 12467–12472 (1996).

21. C. S. Hsieh, S. E. Macatonia, A. O’Garra, K. M. Murphy, T cell genetic background determines default t helper phenotype development in vitro. J. Exp. Med. 181, 713–721 (1995).

22. M. R. Howitt, Y. G. Cao, M. B. Gologorsky, J. A. Li, A. L. Haber, M. Biton, J. Lang, M. Michaud, A. Regev, W. S. Garrett, The Taste Receptor TAS1R3 Regulates Small Intestinal Tuft Cell Homeostasis. ImmunoHorizons 4, 23–32 (2020).

23. S. Laffont, E. Blanquart, M. Savignac, C. Cénac, G. Laverny, D. Metzger, J. P. Girard, G. T. Belz, L. Pelletier, C. Seillet, J. C. Guéry, Androgen signaling negatively controls group 2 innate lymphoid cells. J. Exp. Med. 214, 1581–1592 (2017).

24. L. Mathä, H. Shim, C. A. Steer, Y. H. Yin, I. Martinez, G. Fumio Takei, Female and male mouse lung group 2 innate lymphoid cells differ in gene expression profiles and cytokine production. PLoS One 14(2019), doi:10.1371/journal.pone.0214286.

25. J. Y. Cephus, M. T. Stier, H. Fuseini, J. A. Yung, S. Toki, M. H. Bloodworth, W. Zhou, K. Goleniewska, J. Zhang, S. L. Garon, R. G. Hamilton, V. V. Poloshukin, K. L. Boyd, R. S. Peebles, D. C. Newcomb, Testosterone Attenuates Group 2 Innate Lymphoid Cell-Mediated Airway Inflammation. Cell Rep. 21, 2487–2499 (2017).

26. C. Wang, Z. Bin Xu, Y. Q. Peng, H. Y. Zhang, Q. N. Yu, Y. B. Guo, W. P. Tan, Y. L. Liu, X. C. Meng, S. Bin Fang, D. Chen, Q. L. Fu, Sex differences in group 2 innate lymphoid cell-dominant allergic airway inflammation. Mol. Immunol. 128, 89–97 (2020).

27. S. Kadel, E. Ainsua-Enrich, I. Hatipoglu, S. Turner, S. Singh, S. Khan, S. Kovats, A Major Population of Functional KLRG1 –ILC2s in Female Lungs Contributes to a Sex Bias in ILC2 Numbers. ImmunoHorizons 2, 74–86 (2018).

28. S. Picelli, Å. K. Björklund, B. Reinius, S. Sagasser, G. Winberg, R. Sandberg, Tn5 transposase and tagmentation procedures for massively scaled sequencing projects. Genome Res. 24, 2033–2040 (2014).

29. X. S. Wu, X. Y. He, J. J. Ipsaro, Y. H. Huang, J. B. Preall, D. Ng, Y. T. Shue, J. Sage, M. Egeblad, L. Joshua-Tor, C. R. Vakoc, OCA-T1 and OCA-T2 are coactivators of POU2F3 in the tuft cell lineage. Nature 607, 169–175 (2022).

30. A. P. Szczepanski, N. Tsuboyama, J. Watanabe, R. Hashizume, Z. Zhao, L. Wang, POU2AF2/C11orf53 functions as a coactivator of POU2F3 by maintaining chromatin accessibility and enhancer activity. Sci. Adv. 8, 2403 (2022).

31. D. B. Schubart, A. Rolink, M. H. Kosco-Vilbois, F. Botteri, P. Matthias, B-cell-specific coactivator OBF-1/OCA-B/Bob1 required for immune response and germinal centre formation. Nature 383, 538–542 (1996).

32. D. Chasman, K. Cepek, P. A. Sharp, C. O. Pabo, Crystal structure of an OCA-B peptide bound to an Oct-1 POU domain/octamer DNA complex: specific recognition of a protein-DNA interface. (1999) (available at www.genesdev.org).

33. J. M. Behnke, J. M. Mugambi, S. Clifford, F. A. Iraqi, R. L. Baker, J. P. Gibson, D. Wakelin, Genetic variation in resistance to repeated infections with Heligmosomoides polygyrus bakeri, in inbred mouse strains selected for the mouse genome project. Parasite Immunol. 28, 85–94 (2006).

34. W. Wojciechowski, D. P. Harris, F. Sprague, B. Mousseau, M. Makris, K. Kusser, T. Honjo, K. Mohrs, M. Mohrs, T. Randall, F. E. Lund, Cytokine-Producing Effector B Cells Regulate Type 2 Immunity to H. polygyrus. Immunity 30, 421–433 (2009).

35. R. M. Anthony, J. F. Urban, F. Alem, H. A. Hamed, C. T. Rozo, J. L. Boucher, N. Van Rooijen, W. C. Gause, Memory TH2 cells induce alternatively activated macrophages to mediate protection against nematode parasites. Nat. Med. 2006 128 12, 955–960 (2006).

36. C. B. Wilen, S. Lee, L. L. Hsieh, R. C. Orchard, C. Desai, B. L. Hykes, M. R. McAllaster, D. R. Balce, T. Feehley, J. R. Brestoff, C. A. Hickey, C. C. Yokoyama, Y.-T. Wang, D. A. MacDuff, D. Kreamalmayer, M. R. Howitt, J. A. Neil, K. Cadwell, P. M. Allen, S. A. Handley, M. van Lookeren Campagne, M. T. Baldridge, H. W. Virgin, Tropism for tuft cells determines immune promotion of norovirus pathogenesis. Science 360, 204–208 (2018).

37. Y. H. Huang, O. Klingbeil, X. Y. He, X. S. Wu, G. Arun, B. Lu, T. D. D. Somerville, J. P. Milazzo, J. E. Wilkinson, O. E. Demerdash, D. L. Spector, M. Egeblad, J. Shi, C. R. Vakoc, POU2F3 is a master regulator of a tuft cell-like variant of small cell lung cancer. Genes Dev. 32, 915–928 (2018).

38. N. Goto, A. Fukuda, Y. Yamaga, T. Yoshikawa, T. Maruno, H. Maekawa, S. Inamoto, K. Kawada, Y. Sakai, H. Miyoshi, M. M. Taketo, T. Chiba, H. Seno, Lineage tracing and targeting of IL17RB+ tuft cell-like human colorectal cancer stem cells. Proc. Natl. Acad. Sci. U. S. A. 116, 12996–13005 (2019).

39. A. Tenesa, S. M. Farrington, J. G. D. Prendergast, M. E. Porteous, M. Walker, N. Haq, R. A. Barnetson, E. Theodoratou, R. Cetnarskyj, N. Cartwright, C. Semple, A. J. Clark, F. J. L. Reid, L. A. Smith, K. Kavoussanakis, T. Koessler, P. D. P. Pharoah, S. Buch, C. Schafmayer, J. Tepel, S. Schreiber, H. Völzke, C. O. Schmidt, J. Hampe, J. Chang-Claude, M. Hoffmeister, H. Brenner, S. Wilkening, F. Canzian, G. Capella, V. Moreno, I. J. Deary, J. M. Starr, I. P. M. Tomlinson, Z. Kemp, K. Howarth, L. Carvajal-Carmona, E. Webb, P. Broderick, J. Vijayakrishnan, R. S. Houlston, G. Rennert, D. Ballinger, L. Rozek, S. B. Gruber, K. Matsuda, T. Kidokoro, Y. Nakamura, B. W. Zanke, C. M. T. Greenwood, J. Rangrej, R. Kustra, A. Montpetit, T. J. Hudson, S. Gallinger, H. Campbell, M. G. Dunlop, Genome-wide association scan identifies a colorectal cancer susceptibility locus on 11q23 and replicates risk loci at 8q24 and 18q21. Nat. Genet. 40, 631 (2008).

40. B. T. Harris, V. Rajasekaran, J. P. Blackmur, A. O’Callaghan, K. Donnelly, M. Timofeeva, P. G. Vaughan-Shaw, F. V. N. Din, M. G. Dunlop, S. M. Farrington, Transcriptional dynamics of colorectal cancer risk associated variation at 11q23.1 correlate with tuft cell abundance and marker expression in silico. Sci. Reports 2022 121 12, 1–13 (2022).

41. F. Gerbe, J. H. Van Es, L. Makrini, B. Brulin, G. Mellitzer, S. Robine, B. Romagnolo, N. F. Shroyer, J. F. Bourgaux, C. Pignodel, H. Clevers, P. Jay, Distinct ATOH1 and Neurog3 requirements define tuft cells as a new secretory cell type in the intestinal epithelium. J. Cell Biol. 192, 767–780 (2011).

42. A. D. Gracz, L. A. Samsa, M. J. Fordham, D. C. Trotier, B. Zwarycz, Y. H. Lo, K. Bao, J. Starmer, J. R. Raab, N. F. Shroyer, R. L. Reinhardt, S. T. Magness, Sox4 Promotes Atoh1-Independent Intestinal Secretory Differentiation Toward Tuft and Enteroendocrine Fates. Gastroenterology 155, 1508–1523.e10 (2018).

43. C. A. Herring, A. Banerjee, E. T. McKinley, A. J. Simmons, J. Ping, J. T. Roland, J. L. Franklin, Q. Liu, M. J. Gerdes, R. J. Coffey, K. S. Lau, Unsupervised Trajectory Analysis of Single-Cell RNA-Seq and Imaging Data Reveals Alternative Tuft Cell Origins in the Gut. Cell Syst. 6, 37–51.e9 (2018).

44. M. Bjerknes, C. Khandanpour, T. Möröy, T. Fujiyama, M. Hoshino, T. J. Klisch, Q. Ding, L. Gan, J. Wang, M. G. Martín, H. Cheng, Origin of the brush cell lineage in the mouse intestinal epithelium. Dev. Biol. 362, 194–218 (2012).

45. P. Desai, H. Janova, J. P. White, T. S. Stappenbeck, L. B. Thackray, M. S. Diamond Correspondence, G. V Reynoso, H. D. Hickman, M. T. Baldridge, J. F. Urban, M. S. Diamond, Enteric helminth coinfection enhances host susceptibility to neurotropic flaviviruses via a tuft cell-IL-4 receptor signaling axis. Cell 184(2021), doi:10.1016/j.cell.2021.01.051.

46. A. Banerjee, C. A. Herring, B. Chen, H. Kim, A. J. Simmons, A. N. Southard-Smith, M. M. Allaman, J. R. White, M. C. Macedonia, E. T. Mckinley, M. A. Ramirez Solano, E. A. Scoville, Q. Liu, K. T. Wilson, R. J. Coffey, M. K. Washington, J. A. Goettel, K. S. Lau, Succinate Produced by Intestinal Microbes Promotes Specification of Tuft Cells to Suppress Ileal Inflammation. Gastroenterology 0(2020), doi:10.1053/j.gastro.2020.08.029.

47. G. Krasteva, B. J. Canning, P. Hartmann, T. Z. Veres, T. Papadakis, C. Mühlfeld, K. Schliecker, Y. N. Tallini, A. Braun, H. Hackstein, N. Baal, E. Weihe, B. Schütz, M. Kotlikoff, I. Ibanez-Tallon, W. Kummer, Cholinergic chemosensory cells in the trachea regulate breathing. Proc. Natl. Acad. Sci. U. S. A. 108, 9478–9483 (2011).

48. A. Perniss, S. Liu, B. Boonen, M. Keshavarz, A. L. Ruppert, T. Timm, U. Pfeil, A. Soultanova, S. Kusumakshi, L. Delventhal, Ö. Aydin, M. Pyrski, K. Deckmann, T. Hain, N. Schmidt, C. Ewers, A. Günther, G. Lochnit, V. Chubanov, T. Gudermann, J. Oberwinkler, J. Klein, K. Mikoshiba, T. Leinders-Zufall, S. Offermanns, B. Schütz, U. Boehm, F. Zufall, B. Bufe, W. Kummer, Chemosensory Cell-Derived Acetylcholine Drives Tracheal Mucociliary Clearance in Response to Virulence-Associated Formyl Peptides. Immunity 52, 683–699.e11 (2020).

49. M. I. Hollenhorst, I. Jurastow, R. Nandigama, S. Appenzeller, L. Li, J. Vogel, S. Wiederhold, M. Althaus, M. Empting, J. Altmüller, A. K. H. Hirsch, V. Flockerzi, B. J. Canning, A. E. Saliba, G. Krasteva-Christ, Tracheal brush cells release acetylcholine in response to bitter tastants for paracrine and autocrine signaling. FASEB J. 34, 316–332 (2020).

50. M. Keshavarz, S. F. Tabrizi, A.-L. Ruppert, U. Pfeil, Y. Schreiber, J. Klein, I. Brandenburger, G. Lochnit, S. Bhushan, A. Perniss, K. Deckmann, P. Hartmann, M. Meiners, P. Mermer, A. Rafiq, S. Winterberg, T. Papadakis, D. Thomas, C. Angioni, J. Oberwinkler, V. Chubanov, T. Gudermann, U. Gärtner, S. Offermanns, B. Schütz, W. Kummer, Cysteinyl leukotrienes and acetylcholine are biliary tuft cell cotransmitters. Sci. Immunol. 7, 6734 (2022).

51. C. E. O’Leary, J. Sbierski-Kind, M. E. Kotas, J. C. Wagner, H.-E. Liang, A. W. Schroeder, J. C. de Tenorio, J. von Moltke, R. R. Ricardo-Gonzalez, W. L. Eckalbar, A. B. Molofsky, C. Schneider, R. M. Locksley, Bile acid–sensitive tuft cells regulate biliary neutrophil influx. Sci. Immunol. 7, 1080 (2022).

52. R. Corbett-Detig, R. Nielsen, A Hidden Markov Model Approach for Simultaneously Estimating Local Ancestry and Admixture Time Using Next Generation Sequence Data in Samples of Arbitrary Ploidy. PLoS Genet. 13(2017), doi:10.1371/journal.pgen.1006529.

53. C. J. C. Johnston, E. Robertson, Y. Harcus, J. R. Grainger, G. Coakley, D. J. Smyth, H. J. Mcsorley, R. Maizels, C.: Johnston, C. J. Robertson, J. R. Coakley, G. Smyth, D. J. Mcsorley, Cultivation of Heligmosomoides Polygyrus: An Immunomodulatory Nematode Parasite and its Secreted Products. J. Vis. Exp, 52412 (2015).

54. D. Voehringer, T. A. Reese, X. Huang, K. Shinkai, R. M. Locksley, Type 2 immunity is controlled by IL-4/IL-13 expression in hematopoietic non-eosinophil cells of the innate immune system. J. Exp. Med. 203, 1435–1446 (2006).

55. S. Steinfelder, S. Rausch, D. Michael, A. A. Kühl, S. Hartmann, Immunity to infection Intestinal helminth infection induces highly functional resident memory CD4 + T cells in mice. Eur. J. Immunol 47, 353–363 (2017).

56. D. W. Strong, L. B. Thackray, T. J. Smith, H. W. Virgin, Protruding Domain of Capsid Protein Is Necessary and Sufficient To Determine Murine Norovirus Replication and Pathogenesis In Vivo. J. Virol. 86, 2950–2958 (2012).

57. K. A. Chachu, A. D. LoBue, D. W. Strong, R. S. Baric, H. W. Virgin, Immune Mechanisms Responsible for Vaccination against and Clearance of Mucosal and Lymphatic Norovirus Infection. PLOS Pathog. 4, e1000236 (2008).

58. M. T. Baldridge, S. Lee, J. J. Brown, N. McAllister, K. Urbanek, T. S. Dermody, T. J. Nice, H. W. Virgin, Expression of Ifnlr1 on Intestinal Epithelial Cells Is Critical to the Antiviral Effects of Interferon Lambda against Norovirus and Reovirus. J. Virol. 91, 2079–2095 (2017).

59. L. Baert, C. E. Wobus, E. Van Coillie, L. B. Thackray, J. Debevere, M. Uyttendaele, Detection of murine norovirus 1 by using plaque assay, transfection assay, and real-time reverse transcription-PCR before and after heat exposure. Appl. Environ. Microbiol. 74, 543–546 (2008).

60. M. T. Baldridge, T. J. Nice, B. T. McCune, C. C. Yokoyama, A. Kambal, M. Wheadon, M. S. Diamond, Y. Ivanova, M. Artyomov, H. W. Virgin, Commensal microbes and interferon-λ determine persistence of enteric murine norovirus infection. Science (80-.). 347, 266–269 (2015).

61. T. Sato, H. Clevers, Primary mouse small intestinal epithelial cell cultures. Methods Mol. Biol. 945, 319–328 (2013).

