## Supplementary figures and images for "Genetic mapping reveals *Pou2af2*-dependent tuning of tuft cell differentiation and intestinal type 2 immunity"

### Supplemental Fig 1

#### Supplemental Figure 1

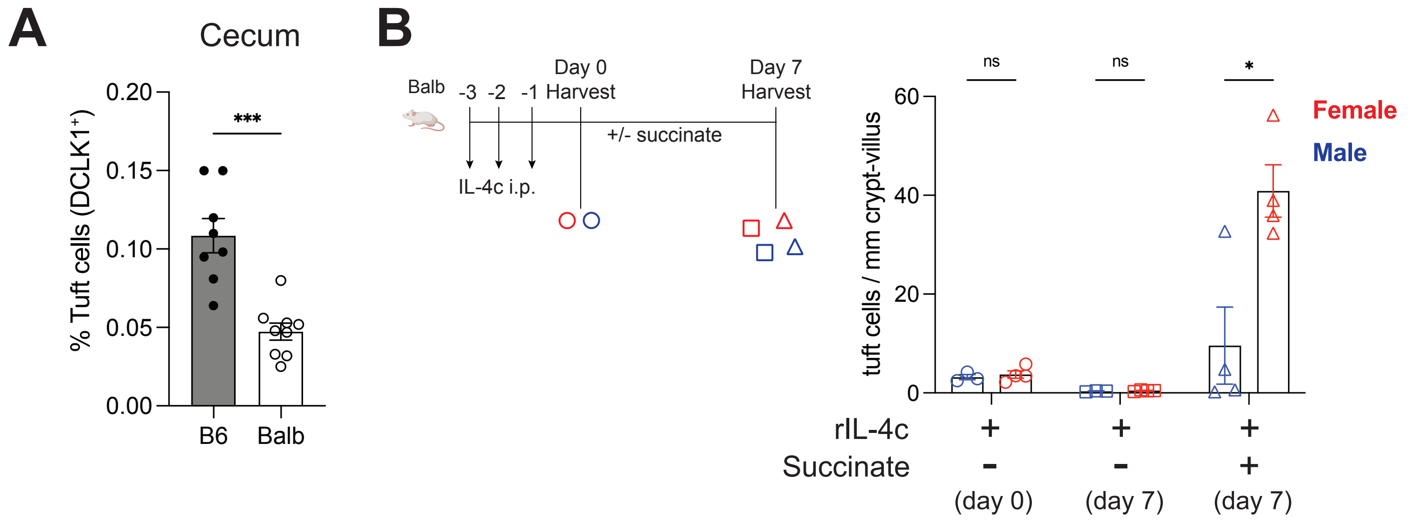

### Supplemental Fig 2

#### Supplemental Figure 2

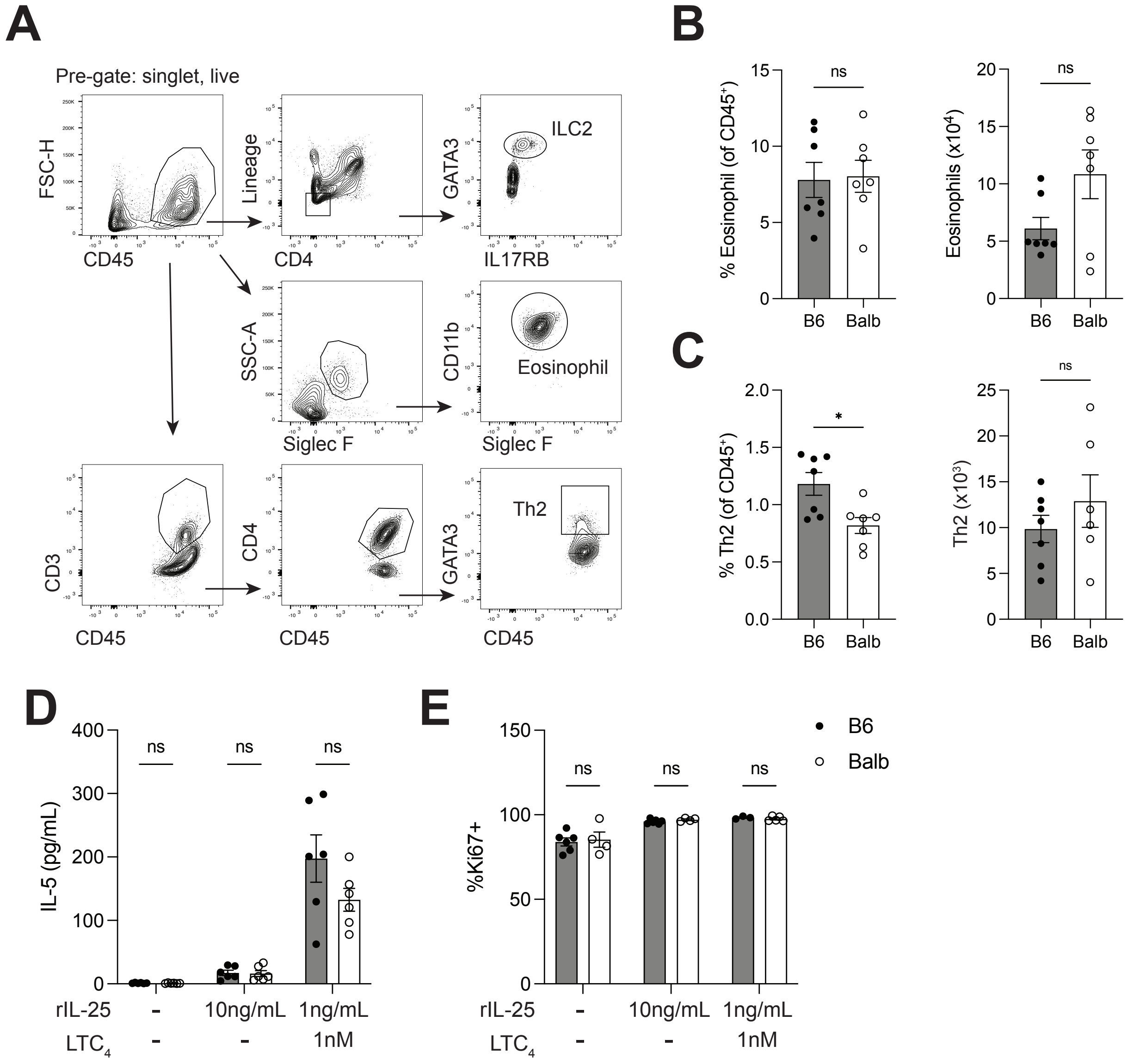

### Supplemental Fig 3

#### Supplemental Figure 3

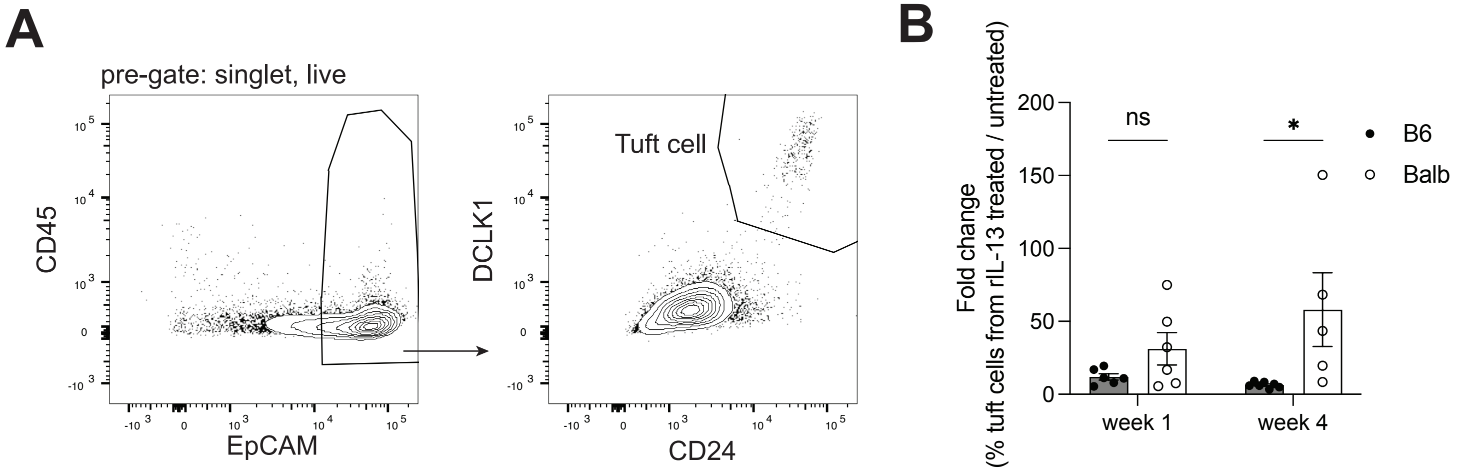

### Supplemental Fig 4

#### Supplemental Figure 4

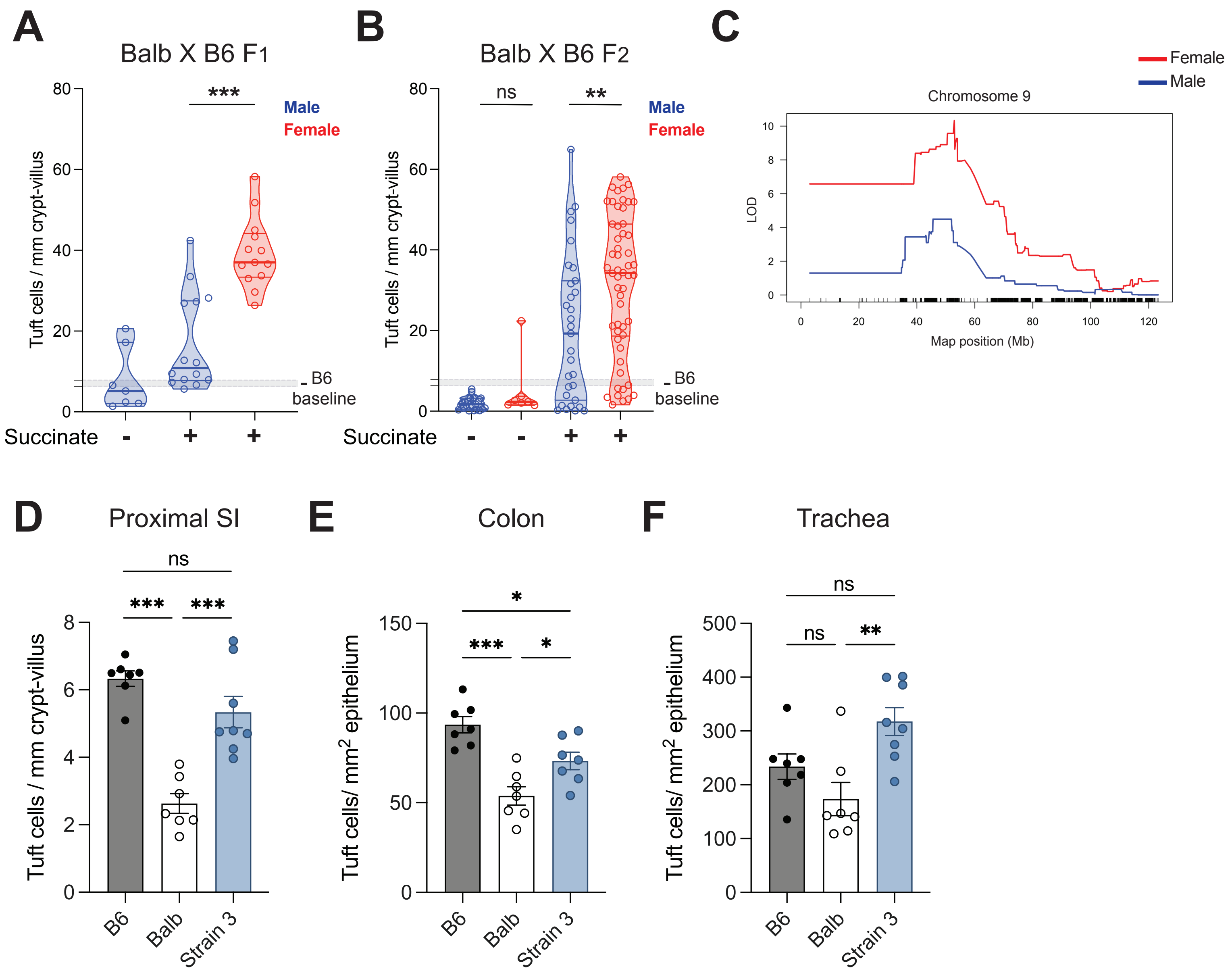

### Supplemental Fig 5

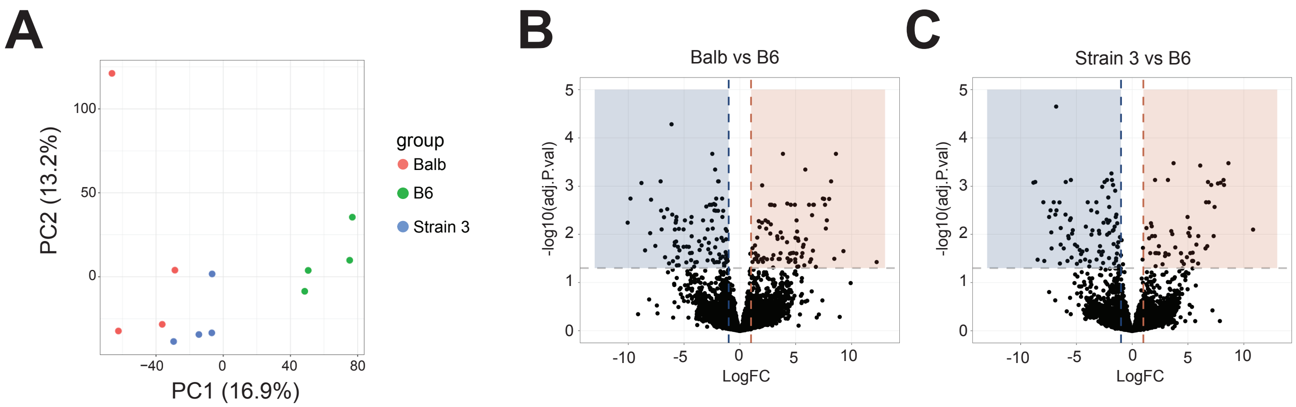

### Supplemental Fig 6

#### Supplemental Figure 6

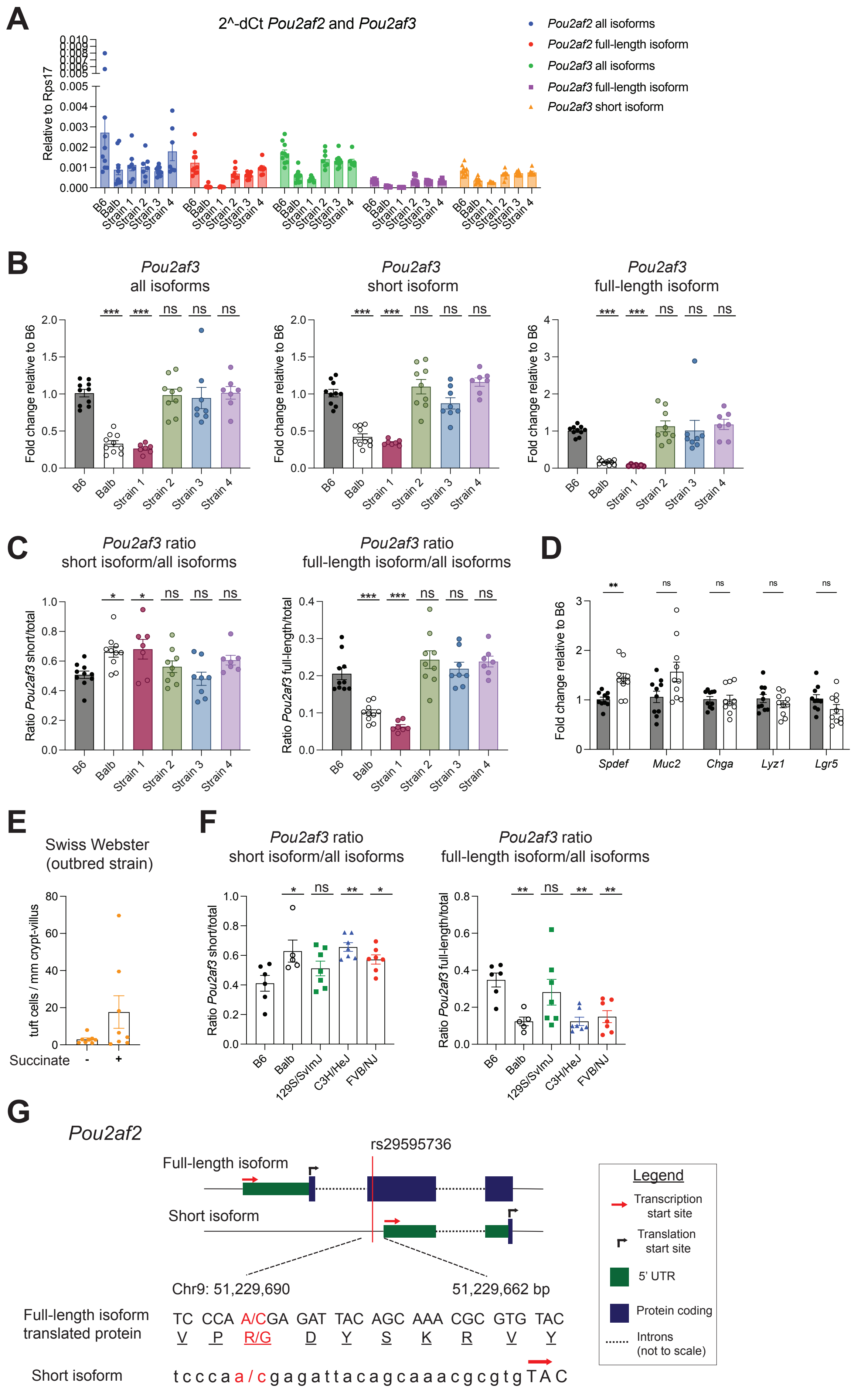

### Supplemental Fig 7

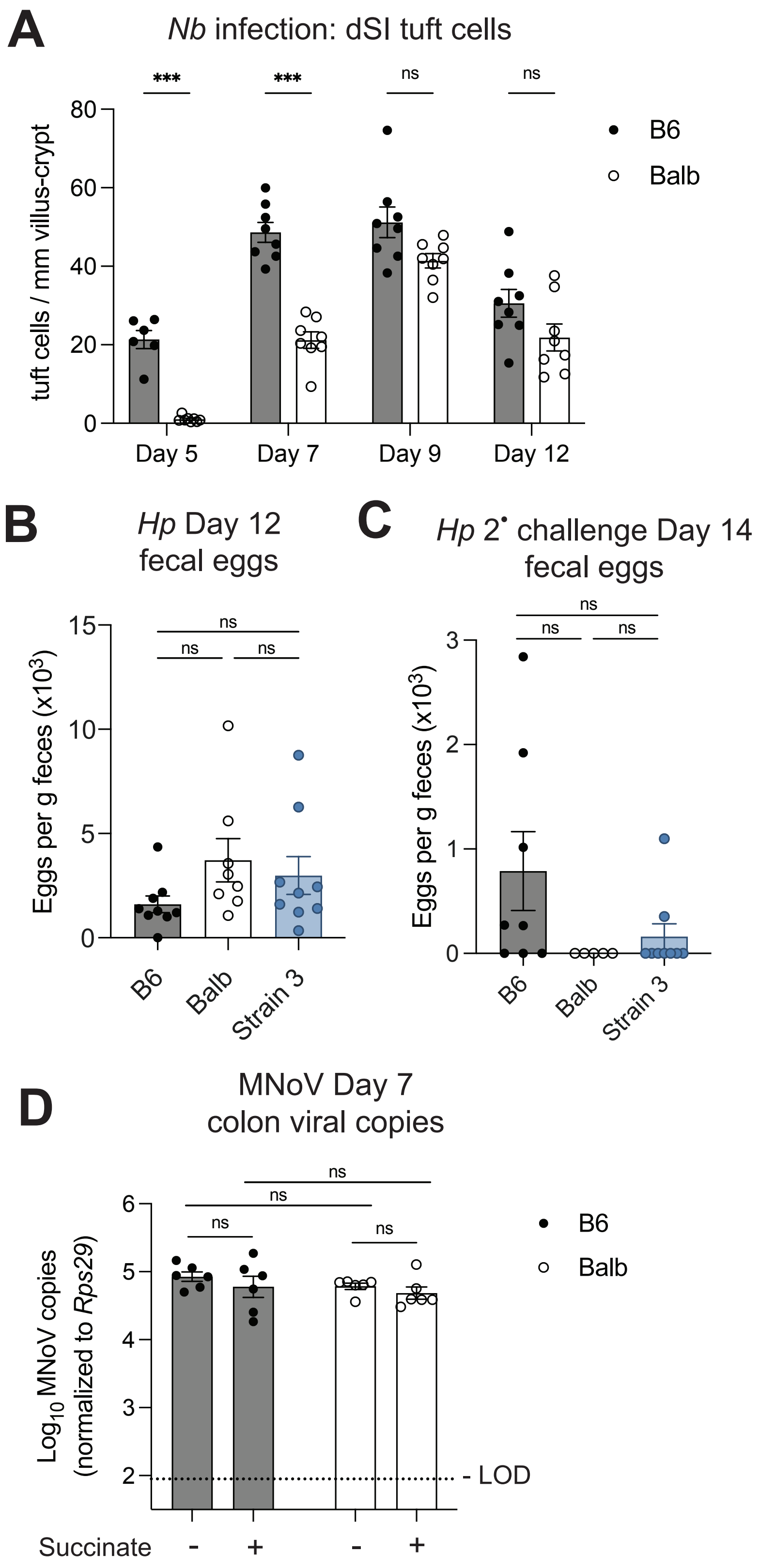
